# Chikungunya Virus Infection of Monocytes Generates a Persistent Macrophage Reservoir That Promotes IFN-π/IL-27–STAT1- and NF-κB–Driven Chronic Arthritis

**DOI:** 10.64898/2026.09.24.754283

**Authors:** Juan Felipe Valdés-López, Y.S Tamayo-Molina, Lady Johana Hernández-Sarmiento, Silvio Urcuqui-Inchima

## Abstract

Chikungunya virus (CHIKV) infection causes acute febrile illness that frequently progresses to chronic arthralgia. Although monocytes are CHIKV targets, whether viral exposure drives their differentiation into macrophages to sustain persistent inflammation remains incompletely understood. To address this, we integrated transcriptomic profiling of pediatric CHIKV patients during acute and convalescent phases, *in vitro* differentiation of primary human monocytes into macrophages under CHIKV infection, and single-cell RNA-sequencing (scRNA-seq) of joint-associated macrophages from an immunocompetent mouse model of chronic CHIKV arthritis at 28 days post-infection.

In acute patients, we observed a marked upregulation of monocyte/macrophage markers alongside robust, IFN-dependent JAK-STAT signaling. *In vitro*, CHIKV-infected monocytes differentiated into persistently infected macrophages (MDM-CHIKV) that closely aligned with an M1-polarized transcriptional profile. These cells exhibited STAT1/NF-κB-dependent inflammation, ISG induction, and enhanced antigen presentation capacity. Remarkably, MDM-CHIKV uniquely produced IFN-π/IL-27, revealing an alternative non-canonical axis where JAK-STAT activation persists despite the absence of classical type I IFNs. Functionally, MDM-CHIKV displayed an enhanced respiratory burst upon immune complex uptake, pointing to FcγR-mediated pathology, as well as complement activation and T-cell stimulatory capacity. However, upon TLR4 challenge, these cells retained partial regulatory responses via IL-10 and TGF-β; this hybrid immunophenotype offers a potential mechanistic explanation for relapsing arthralgia. Additionally, scRNA-seq of murine joints confirmed that persistent CHIKV replication was restricted to a distinct macrophage subset that resembles MDM-CHIKV, establishing these cells as the primary *in vivo* viral reservoir. Mirroring our human findings, joint-associated murine macrophage subsets exhibited elevated *IFN-π*/*IL-27* expression alongside robust NF-κB/STAT1-dependent inflammatory signatures (*Tnf*, *Il1β*, Il6, *Ccl2*, *Ccl5*, *Cxcl9*, *Cxcl10*, *Cxcl1–3,* and/or *Ptgs2*), underscoring their critical role in sustaining chronic arthritis.

Collectively, our findings demonstrate that CHIKV actively drives monocyte-to-macrophage differentiation into a persistently infected, M1-skewed hybrid phenotype that serves as a viral reservoir and fuels pro-arthritic inflammation. Consequently, macrophage reprogramming, the non-canonical IFN-π/IL-27–STAT1 axis, FcγR-mediated phagocytosis, and complement activation emerge as central pathogenic nodes and promising therapeutic targets for mitigating chronic post-CHIKV arthropathy.

**Author Summary:** Chikungunya virus (CHIKV) is a mosquito-borne pathogen that causes acute fever and, in a substantial proportion of patients, progresses to chronic, debilitating joint pain that can last for months or even years. The mechanisms by which the virus establishes this persistent arthritic disease remain poorly understood, and effective treatments for chronic CHIKV arthritis are lacking.

In this study, we investigated whether CHIKV infection of monocytes, immune cells that circulate in the blood, actively drives their transformation into macrophages, a cell type that resides in tissues and plays a central role in inflammation. By analyzing blood samples from children with acute and convalescent CHIKV disease, we found strong signs that monocyte-to-macrophage differentiation is actively occurring during natural infection. Using an *in vitro* model, we showed that CHIKV-infected monocytes differentiate into macrophages that become persistently infected, carrying the virus for extended periods. These virus-exposed macrophages adopt a mixed activation state, combining features of classical pro-inflammatory (M1) and regulatory (M2) macrophages. Notably, they produce a cytokine called IFN-π/IL-27, which drives chronic inflammation through a specific signaling pathway, while also showing enhanced capacity to activate T cells, trigger complement cascades, and mount strong oxidative responses upon encountering immune complexes, features that may explain the waxing and waning joint symptoms seen in patients.

To confirm these findings in a living system, we analyzed joint tissues from a mouse model of chronic CHIKV arthritis using single-cell technology. We discovered that a specific subset of joint-resident macrophages serves as the primary viral reservoir, actively replicating the virus and producing IFN-π/IL-27 and other inflammatory molecules that perpetuate joint inflammation and damage.

Collectively, our work reveals that CHIKV hijacks the normal differentiation process of monocytes to create a long-lived, virus-harboring macrophage population that fuels chronic joint inflammation. We identify the IFN-π/IL-27 signaling pathway, complement activation, and antibody-driven macrophage responses as key drivers of disease. These findings open new avenues for therapeutic intervention, suggesting that targeting these pathways could alleviate chronic arthritis in patients suffering from long-term CHIKV disease.

## 1. Introduction

Chikungunya virus (CHIKV) is an arthropod-borne, positive-sense single-stranded RNA (ssRNA+) virus transmitted to humans through the bite of infected female *Aedes* mosquitoes (1)(2). It is the etiological agent of Chikungunya disease (CHIKD), an acute febrile illness characterized by severe musculoskeletal inflammation and prominent joint involvement (1)(2). Although the acute phase typically resolves within 1–2 weeks, a substantial proportion of patients develop chronic symptoms, including persistent arthritis, tenosynovitis, and debilitating arthralgia that can last for months to years after viral clearance from the bloodstream (3)(4). This chronic phase is thought to be driven by immunopathological mechanisms associated with elevated levels of pro-inflammatory mediators, including tumor necrosis factor alpha (TNF-α), Interleukin 1 beta (IL-1β), IL-6, IL-8, IL-12, IL-15, and selected chemokines such as CCL-2, CCL-5, CCL-8, CXCL-9/10/11, which have been detected in both patient sera and *in vitro* CHIKV-infected monocytes and macrophages (5)(6)(7). Several of these mediators correlate with disease severity and viral load, highlighting their potential role in disease progression (8)(6,9)(10). Intriguingly, elevated levels of IL-27, recently proposed as a putative type V interferon-pi (IFN-π) (11)(12), have also been reported in patients with chronic CHIKD, where they correlate positively with the number of painful joints (13). Furthermore, IFN-π/IL-27 is abundantly produced in supernatants from CHIKV-infected macrophages *in vitro* (14,15), suggesting a potential role for this cytokine in the maintenance of arthralgia.

Monocytes and macrophages are innate mononuclear phagocytes that play critical roles in orchestrating antiviral defenses and regulating inflammation (16)(17)(18). As precursors of tissue macrophages, circulating monocytes undergo complex differentiation upon recruitment into tissues, a process shaped by microenvironmental cues, including pathogen- and damage-associated molecular patterns (PAMPs and DAMPs, respectively), cytokines, and growth factors (19)(20). This transition involves distinct morphological changes, including increased cell size, enhanced cytoplasmic complexity, and a reduced nuclear-cytoplasmic ratio (21). These structural shifts are accompanied by dynamic regulation of pattern recognition receptors (PRRs), such as Toll-like receptors (TLRs), as well as phagocytic receptors including C-type lectins, Fcγ receptors, and scavenger receptors (22)(21)(23). Once differentiated, macrophages acquire long-lived phenotypes that enable tissue surveillance, clearance of pathogens and apoptotic cells, and maintenance of tissue homeostasis (19)(24). A defining feature of macrophage biology is their ability to polarize into specialized functional states in response to local signals. Classically activated (M1) macrophages arise under Th1-type stimulation (e.g., IFN-γ + TNF-α or LPS) and mount pro-inflammatory, antimicrobial responses through NF-κB and STAT1-dependent production of cytokines, chemokines, and reactive oxygen species (25)(26). These cells are characterized by high expression of TLRs, costimulatory molecules, and MHC class II, thereby promoting Th1 responses and enhanced phagocytic capacity (27)(28)(25). In contrast, alternatively activated (M2) macrophages arise in response to distinct microenvironmental cues. While the Th2 cytokines IL-4 and IL-13 drive M2a polarization, immune complexes combined with TLR or IL-1R agonists promote the M2b phenotype, and IL-10 or glucocorticoids induce M2c differentiation. Additionally, tumor microenvironment signals generate M2d macrophages, commonly known as tumor-associated macrophages. These M2 subsets generally facilitate the resolution of inflammation and promote tissue repair, predominantly through STAT6- and/or STAT3-mediated signaling pathways (27)(28)(25). M2 macrophages upregulate receptors such as the macrophage mannose receptor (CD206), Dectin-1, CD163, and CD36, and produce anti-inflammatory cytokines (IL-10), pro-fibrotic mediators (arginase-1, TGF-β), and chemokines (CCL-17, CCL-18) (27)(28)(25)(29). This polarization spectrum, encompassing multiple intermediate states, positions macrophages as central players of both host defense and immunopathology during viral infections.

During CHIKV infection, musculoskeletal and joint tissues in both patients and experimental models exhibit heavy infiltration by monocytes and macrophages. Several studies suggest that these cells contribute to both viral control and disease pathogenesis during acute and chronic phases of infection (30)(31)(14,23). Additionally, macrophages have been proposed to serve as viral reservoirs in persistent infections, as observed in non-human primate and mouse models (32)(33).

Monocytes have emerged as central players in CHIKV pathogenesis. Her et al. (2010) first established that monocytes from patients with acute CHIKV infection constitute the primary targets of CHIKV infection in peripheral blood and further demonstrated their susceptibility to productive infection *in vitro* (34). Extending these observations, Michlmayr et al. (2018) reported that CHIKV infection drives classical monocytes (CD14⁺⁺CD16⁻, CM) toward an intermediate phenotype (CD14⁺⁺CD16⁺, IM). Furthermore, the detection of CHIKV-E1 antigen in IM from pediatric patients (35) suggests that these cells support viral replication and may act as a “Trojan horse” facilitating tissue dissemination (31). In line with this, our previous work confirmed that human monocytes are permissive to CHIKV infection *in vitro* (23)(36), and demonstrated that infection induces the expansion of both IM and non-classical (CD14⁺CD16⁺⁺, NM) monocyte subsets, even in the absence of other peripheral blood mononuclear cell (PBMC) populations (37). This subset shift was accompanied by transcriptional upregulation of antiviral PRR pathways. Notably, CHIKV did not activate canonical NF-κB signaling, but instead triggered robust type I interferon (IFN-I)-dependent STAT1-driven inflammatory and antiviral transcriptional programs. Through pharmacological inhibition, we identified TLR3, TLR8, and RIG-I, but not TLR2 or TLR4, as essential mediators of monocyte subset differentiation, with TLR3 and RIG-I additionally governing IFN-I-mediated antiviral responses, showing that CHIKV-RNAs are the main drivers of monocyte activation and differentiation (37).

Together, these findings identify human monocytes as key targets of CHIKV infection, both *in vitro* and *in vivo,* and as capable of mounting strong pro-inflammatory and antiviral responses. Despite this evidence, the impact of CHIKV infection on monocyte-to-macrophage differentiation and its role in chikungunya immunopathogenesis remain poorly understood. Because macrophages are pivotal orchestrators of inflammation, tissue repair, and viral persistence, elucidating how CHIKV skews this differentiation process is critical to unlocking the mechanisms driving chronic disease. Therefore, this study aimed to characterize the morphological, phenotypic, and functional alterations in human monocytes differentiated into macrophages in the presence or absence of CHIKV. By elucidating how CHIKV shapes macrophage lineage commitment and polarization, these findings advance our understanding of viral immunopathogenesis and highlight potential targets for therapeutic intervention in chronic CHIKD.

## 2. Materials and methods Ethics statement

The Human Ethics Committee of the “Sede de Investigación Universitaria, Universidad de Antioquia” approved the study (April 26, 2022; minutes No. 028). All individuals who voluntarily participated in this study provided written informed consent, in accordance with the principles of the Declaration of Helsinki. Four healthy donors were included in this study.

### 2.1. Viral stocks and CHIKV titration

A Colombian clinical isolate of CHIKV obtained following the protocol described in (38), kindly gifted by Professor Francisco Javier Díaz (University of Antioquia), was amplified in Vero cells (ATCC CCL-81), as previously reported (14,23). Briefly, cells were grown in Dulbecco’s Modified Eagle Medium (DMEM; Sigma-Aldrich, St. Louis, USA) supplemented with 5% heat-inactivated fetal bovine serum (FBS; Gibco, Thermo Fisher Scientific, Massachusetts, USA), 4 mM L-glutamine (Sigma-Aldrich), 0.3% (v/v) sodium carbonate (NaCO_3_; Sigma-Aldrich) and 1% (v/v) antibiotic-antimycotic solution (Corning-Cellgro, New York, USA), and incubated at 37°C and 5% CO_2_ in cell culture flasks, to a cell density of 1x10^5^-1x10^6^ cells/mL. Vero cells were inoculated with CHIKV at a multiplicity of infection (MOI) of 0.1. After incubation at 37°C and 5% CO_2_ for 48 hours or until cytopathic effect was observed, supernatants were collected, clarified by centrifugation (1650 x g for 10 min), and stored at -80°C. Viral stocks were titrated by plaque assay on Vero cells, as previously described (14,23).

### 2.2. Isolation of human monocytes and differentiation into macrophages in the presence or absence of CHIKV

Peripheral blood mononuclear cells (PBMCs) from healthy donors were isolated by Lymphoprep density gradient centrifugation (850 × g, 21 min) as previously described (14,23). PBMCs from each donor were processed independently. Platelets were depleted by three washes with PBS-1X (250 × g, 10 min), and CD14⁺ monocytes were quantified by flow cytometry. For monocyte isolation, cells were seeded at 5 × 10⁵ CD14⁺ cells per well in scratched 24-well plates and allowed to adhere for 2 h in RPMI-1640 with 0.5% (v/v) autologous serum or plasma at 37°C with 5% CO₂. To eliminate non-adherent cells, the cultures were gently washed twice with 1X PBS 1X, and adherent monocytes were cultured in complete RPMI-1640 (10% FBS, 1% antibiotic-antimycotic). After 24 h, monocytes were either challenged with CHIKV at an MOI of 10 in serum-free RPMI-1640 for 90 min (23,35) or not. Cells were then washed to remove unbound virus and incubated in complete medium. Both infected and uninfected monocytes were differentiated into unpolarized monocyte-derived macrophages (MDM) and MDM-CHIKV, respectively, by adherence for 5 days in complete medium without exogenous growth factors, with half-medium changes every 48 h.

### 2.3. Flow cytometry analysis

A BD-LSRFortessa flow cytometer (BD Biosciences, New Jersey, USA) was set up to assess cell size (FSC-A) and cytoplasmic complexity/granularity (SSC-A). To evaluated macrophages surface markers expression, MDM and MDM-CHIKV cultures were stained with fluorescently-conjugated anti-human antibodies against CD14 (FITC, clone: 61D1), CD16 (PE, clone: 3G8), CD68 (PE, clone: eBIOY1/82A), CD80 (FITC, clone: 2D10.4), CD86 (PE, clone: FUN-1), CD206 (FITC, clone: 19.2), HLA-DR (FITC, clone: L243), and TLR2 (PE, clone: TLR2.1) (eBioscience, Massachusetts, USA, and BD Bioscience), or corresponding isotype controls. Staining was performed in PBS-1X at 4°C for 40 min in the dark. The cells were washed by centrifugation, resuspended in PBS-1X, and analyzed by flow cytometry. Positive populations were defined by using isotype controls and unstained cells, and compensation was performed to account for spectral overlap. At least 10.000 events were acquired per sample. Data were analyzed using FlowJo software (version 7.6.2), and debris and dead cells were excluded based on forward- and side-light properties.

### 2.5. Microscopy

MDM and MDM-CHIKV cultures were visualized and photographed using an Axio Vert.A1 microscope (ZEISS, Oberkochen, Germany) equipped with a Ds-fi1 camera (Nikon, Tokyo, Japan) and adapted for transmitted-light in-phase contrast and epi-fluorescence microscopy. Images were processed and analyzed using IrfanView (version 4.69), ImageJ (version 1.8.0), and GIMP (version 2.10.30).

### 2.6. RNA extraction, cDNA synthesis, and bulk RNA-Seq of human macrophages

Total RNA was extracted from MDM and MDM-CHIKV cultures using the Direct-zol™ RNA Miniprep Plus kit (Zymo Research, California, USA), following the manufacturer’s protocol. RNA samples were treated with DNase I (Zymo Research) to remove residual genomic DNA, and RNA concentration was measured by Nanodrop spectrophotometry (Thermo Scientific, Massachusetts, USA). For RT-qPCR, 400 ng of total RNA per sample was used for cDNA synthesis using the RevertAid Minus First Strand cDNA Synthesis Kit (Thermo Scientific), according to the manufacturer’s instructions. cDNA samples were stored at -20°C.

For bulk RNA-Seq, 1 μg of total RNA per sample was used for library preparation, and sequencing was performed on an Illumina HiSeq 2000 platform (Macrogen, Seoul, South Korea). Raw image data were converted to FASTQ files, and sequence quality was assessed using FastQC (14). Clean reads were obtained by removing low-quality adapters, poly-N-containing stretches, and reads shorter-than-70 bp. Reads were aligned to the human reference genome (GRCh38) and CHIKV genome (strain LR2006 OPY1) using HISAT2 software (39). Transcript assembly was performed with StringTie (40), and gene-level counts were generated using featureCounts from the Subread software (41). Data will be deposited in the Gene Expression Omnibus (GEO) upon acceptance of the manuscript.

Raw count data were processed in R software (version 4.2.0). Gene annotations, including gene type (e.g., non-coding, pseudogene, or protein-coding), gene symbols, and Entrez IDs, were assigned to each gene. The dataset was filtered to retain only protein-coding genes, and gene lengths were retrieved using the *Homo.sapiens* Library. To identify the top differentially expressed genes (DEGs), we selected genes with a false discovery rate (FDR) < 0.05 and |Log2 Fold Change (FC) > 0.6 (MDM-CHIKV vs MDM), using the DESeq2 library (42). Raw counts were also normalized to transcripts per million (TPM) using R software (version 4.2.0) for visualization and comparative analysis. Based on prior reports (16)(15), gene subsets associated with JAK-STAT signaling, the PRR pathway, and antiviral and inflammatory responses were selected for downstream analysis. Data visualization, including box plots and heatmaps, was performed using GraphPad Prism version 8.0.1 (GraphPad Software Inc., San Diego, CA), and R statistical software.

### 2.7. Transcriptomic analysis of CHIKD pediatric patients

To characterize the transcriptional response of pediatric patients during natural Chikungunya infection, we reanalyzed the publicly available RNA-seq dataset GSE99992 (GEO) generated by Michlmayr et al. (2018). This dataset comprises whole-blood samples collected from 43 pediatric cases at two time points post-symptom onset: acute phase (1–2 days) and convalescent phase (15–17 days). mRNA levels are shown as TPM.

### 2.8. Transcriptomic analysis of M1 and M2 polarized macrophages

To infer the polarization profile of MDM-CHIKV, we reanalyzed the publicly available bulk RNA-seq dataset **GSE205990** from GEO (https://www.ncbi.nlm.nih.gov/geo/query/acc.cgi?acc=GSE205990), generated by Embregts et al. (43). In this dataset, human monocytes from healthy donors (n = 3) were differentiated into MDM in RPMI-1640 supplemented with 10% pooled human serum, 1% v/v GlutaMax, and 20 ng/mL macrophage colony-stimulating factor (M-CSF), for 6 days at 37°C with 5% CO_2_. Then, MDMs were polarized into M1, M2a, and M2c phenotypes by stimulation for 48 h with IFN-γ (20 ng/mL) + LPS (100 ng/mL), IL-4 (20 ng/mL), or IL-10 (20 ng/mL), respectively. To identify the top DEGs, we selected genes with an FDR < 0.05 and |Log2FC |> 0.6 (polarized MDM vs. control MDM), using the DESeq2 package (42).

### 2.9. Validation of RNA-Seq results by RT-qPCR

RT-qPCR was used to confirm the expression of key genes identified by RNA-Seq analysis. First-strand cDNA was generated using SuperScript III reverse transcriptase, and specific primer pairs (Table 1) were used to quantify the mRNA expression of *IL10*, *TGFA*, *TGFB1*, *IL27*, *EBI3*, *SOCS3*, *APOBEC3A*, *ISG15*, *OAS1*, *PKR*, and *ACTB* (14)(15). Brief, PCR amplifications were carried out using the SYBR system (Invitrogen, Oregon, USA). The Bio-Rad CFX Manager was used to obtain the cycle thresholds (Ct) for each sample, determined by fitting a regression to the linear phase of the PCR amplification curve. Relative expression of each target gene was normalized to the unstimulated control and the housekeeping gene *ACTB* (ΔΔCt), and was reported as the Log2 Fold Change.

### 2.10. Activation of the TLR4 signaling pathway in human MDM

Human MDM and MDM-CHIKV were stimulated with 50 ng/mL ultrapure Lipopolysaccharide (LPS, InvivoGen) in fresh complete medium and incubated at 37°C and 5% CO_2_. After 24 h, culture supernatants and cell lysates were collected and stored at -80°C.

### 2.11. Cytokine quantification

Levels of TNF-α, IL-6, CCL-2/MCP-1, and CCL5 were quantified in culture supernatants from MDM and MDM-CHIKV using commercially available ELISA kits (BioLegend) following the manufactureŕs instructions. The detection limit was 2-4 pg/mL. Mature IFN-π/IL-27 levels were measured using the LEGEND MAX™ Human IL27 ELISA Kit (BioLegend) following the manufactureŕs instructions. The detection limit for this assay was 11 pg/mL.

### 2.12. ROS production and phagocytosis assay

To assess reactive oxygen species (ROS) production in response to phagocytosis, MDM were stimulated with Zymosan from *Saccharomyces cerevisiae* (Sigma-Aldrich) or serum-opsonized Zymosan (ZOP). For opsonization, Zymosan (1 mg/mL) was incubated with pooled normal human serum (NHS; n = 3) for 45 min at 37°C, then washed with PBS-1X to remove excess serum, as previously reported (44)(45). MDMs were then stimulated with 10 μg/mL of Zymozan or ZOP for 90 min at 37°C and 5% CO_2_. Cells were subsequently stained with dihydrorhodamine 123 (DHR-123) for 30 min under the same conditions, according to the manufacturer’s instructions. ROS production was measured by flow cytometry as described in section 2.3.

### 2.13. scRNA-seq analysis of joint tissue from mice with chronic CHIKV infection

To explore single-cell transcriptional events associated with chronic CHIKV infection in mice, we reanalyzed the public scRNA-seq dataset GSE300299 from GEO, generated by Zarralla et al (33). In this dataset, wild-type (WT) C57BL/6J mice (n = 3) were inoculated with the CHIKV strain AF15561, and joint-associated tissue from the ipsilateral ankle was harvested at 28 days post-infection (dpi) for 10x Genomics Next GEM Single-Cell 3’ expression profiling. The sparse .h5 matrix containing cell-by-gene read counts was imported into R using the Read10X_h5 function in Seurat (version 5; (46)). Cell type annotation was performed using SingleR with the ImmGen and Mouse RNA-seq reference datasets (47). Data normalization, selection of the top 2,000 highly variable features (via the vst method), data scaling, and principal component analysis (PCA) were performed following the standard Seurat pipeline (46). Data integration across samples was performed using the IntegrateLayers function combined with Harmony, using the first 8 principal components (PCs).

To identify specific macrophage subtypes, we subclustered only those cells annotated as macrophages by the ImmGen and Mouse RNA-seq reference datasets. We then re-ran the Seurat pipeline incorporating Harmony integration across the first 14 PCs, and graph-based clustering was performed at a resolution of 0.2. Differential gene expression analysis of the scRNA-seq data was subsequently conducted using FindMarkers function in Seurat, with genes considered differentially expressed at an adjusted P < 0,05 (46).

### 2.14. Statistical analysis

Statistical analyses and figure generation were performed using R statistical software (R Core Team, 2020). The statistical tests used are indicated in the figure legends. Data are presented as mean ± SD or as median with interquartile range, Log2 FC, or TPM. Statistical significance results were defined as p<0.05 (*).

## 3. Results

### 3.1. CHIKV Drives Monocyte-to-Macrophage Differentiation and Induces a Divergent Innate Immune Response in Acute Chikungunya Disease

During the acute phase of CHIKD, monocytes serve as primary targets of CHIKV infection in the bloodstream, driving type I IFN-dependent antiviral responses while simultaneously facilitating viral replication and tissue dissemination (34)(31)(23)(37). However, how CHIKV infection influences monocyte-to-macrophage differentiation remains poorly understood. To address this gap, we reanalyzed the publicly available RNA-seq dataset GSE99992 (GEO, NCBI) generated by Michlmayr et al. (35) derived from pediatric whole-blood samples during the acute and convalescent phases of CHIKD. Comparing acute and convalescent phases, we identified a significant upregulation of monocytic markers, including *CD14*, *FCGR3A* (CD16A), *CCR1*, *CXCR1*, and the calprotectin subunits, *S100A8* and *S100A9* (Fig 1A). These findings demonstrate robust monocyte activation during acute CHIKV infection *in vivo*.

**Figure 1.**
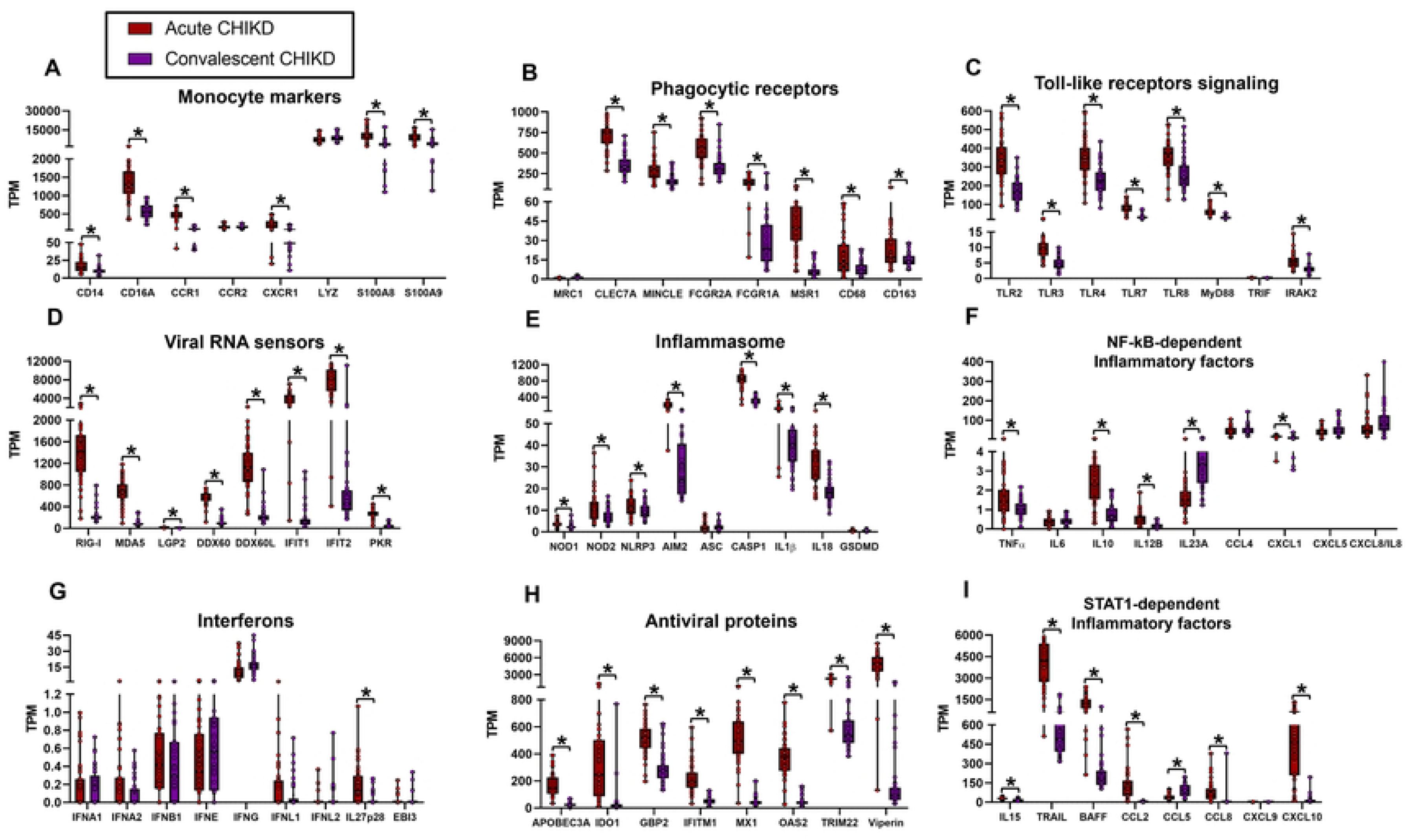
CHIKV infection induces dynamic transcriptional reprogramming of monocyte and innate immune pathways during acute chikungunya disease. Whole-blood mRNA expression levels of key immune gene categories from CHIKD patients during the acute and convalescent phases. A) Monocyte markers. B) Phagocytic receptors. C) Toll-like receptor signaling. D) Viral RNA sensors. E) Inflammasome components. F) NF-κB-dependent inflammatory factors. G) Interferons. H) ISGs encoding antiviral proteins. I) STAT1-dependent inflammatory factors. Data are presented as medians with interquartile range. Normality was assessed using the Shapiro-Wilk test, and sample comparisons were performed using Student’s t-test, whereby * = p < 0.05, comparing acute vs. convalescent CHIKD patient groups.

Consistent with these findings, we also observed significant increases in mRNA levels encoding key macrophage phagocytic receptors during the acute phase of CHIKD (Fig 1B). These included C-type lectin receptors (*CLEC7A/*Dectin-1 and *CLEC4E/*MINCLE), FCγ receptors (*FCGR2A/*CD32A and *FCGR1A*/CD64A), and scavenger receptors (*MSR1*, *CD68*, and *CD163*), suggesting that CHIKV infection promotes monocyte-to-macrophage differentiation *in vivo*.

We also found a significant induction of PRRs involved in CHIKV sensing during the acute phase of CHIKD (Fig 1C). These included cell surface TLRs, such as *TLR2* and *TLR4*, which have been suggested to recognize structural CHIKV PAMPs (15,48), as well as endosomal TLRs (*TLR3*, *TLR7*, and *TLR8*) that sense viral RNA (37,49). This upregulation was accompanied by elevated transcript levels of downstream signaling adaptors, including *MyD88* and IL-1 receptor-associated kinase 2 (*IRAK2*). Collectively, these findings indicate a robust transcriptional activation of the TLR signaling pathway in patients during acute CHIKD.

We detected upregulation of cytoplasmic viral RNA sensors, including retinoic acid-inducible gene I (*DDX58/*RIG-I), melanoma differentiation-associated gene 5 (*IFIH1/*MDA5), and laboratory of genetics and physiology 2 (*DHX58/*LGP2). This was accompanied by elevated transcript levels of DExD/H-Box helicase 60 (*DDX60*), *DDX60L*, interferon-induced protein with tetratricopeptide repeats 1 (*IFIT1*), *IFIT2*, and protein kinase R (*PKR*) (Fig 1D), confirming the induction of a robust antiviral response in patients with acute CHIKD. Furthermore, intracellular PRRs, including the NLRs (*NOD1*, *NOD2*, *NLRP3*) and the DNA sensor *AIM2*, were significantly upregulated, along with inflammasome-related effectors such as Caspase-1 (*CASP1*), *IL1B*, and *IL18* (Fig 1E), suggesting inflammasome priming during acute CHIKV infection.

Interestingly, we observed only a modest induction of classical NF-κB-dependent inflammatory cytokines. Although *TNF*, *IL10*, *IL12B*/IL12p40, and *CXCL1* mRNA levels were significantly increased, other key mediators such as *IL6*, *CCL4*, *CXCL5*, and *CXCL8*/*IL8* showed no significant changes (Fig 1F). In contrast, *IL23A* was selectively upregulated during the convalescent phase. Together, these results suggest that acute inflammation in CHIKD may be partially uncoupled from classical NF-κB signaling.

While transcripts encoding type I (*IFNA1/2*, *IFNB1*, *IFNE*), type II (*IFNG*), and type III (*IFNL1*) interferons were detected in both acute and convalescent patients (Fig 1G), their expression levels did not differ significantly between disease phases. This finding suggests a sustained, concurrent expression of multiple IFN subtypes throughout CHIKV infection. Additionally, we detected significant upregulation of the p28 subunit (*IL27p28*/*IL30*), a component of the IL-27 (IFN-π) heterodimer during the acute phase (Fig. 1G). However, its partner subunit, Epstein-Barr virus-induced gene 3 (*EBI3*), was not co-expressed (Fig 1G), indicating that assembly of the functional IFN-π/IL-27 heterodimer is not consistently induced under these conditions.

Despite the absence of significant differences in systemic IFN levels between acute-phase and convalescent patients, we observed a robust transcriptional upregulation of interferon-stimulated genes (ISGs) encoding antiviral effectors during acute CHIKD. This response was particularly marked in myeloid-lineage-associated transcripts, including Apolipoprotein B mRNA editing enzyme catalytic subunit 3A (*APOBEC3A*) and indoleamine 2,3-dioxygenase 1 (*IDO1*) (Fig 1H), both recognized as key drivers of M1-macrophage differentiation (50)(51). Additionally, we detected strong induction of a broad array of canonical antiviral ISGs, including guanylate binding protein 2 (*GBP2*), interferon-induced transmembrane protein 1 (*IFITM1*), interferon-induced GTP-binding protein (*MX1*), 2’-5’-oligoadenylate synthase 2 (*OAS2*), tripartite motif-containing 22 (*TRIM22*), and Radical S-adenosyl methionine domain-containing protein 2 (*RSAD2/*Viperin), all of which were highly expressed (Fig 1H). We also observed significant upregulation of STAT1-dependent inflammatory factors (11,15), including *IL15*, TNF-related apoptosis-inducing ligand (*TNFSF10/TRAIL*), B-cell activating factor (*TNFSF13B/BAFF*), *CCL2*, *CCL8*, and *CXCL10* during acute CHIKD. In contrast, *CCL5* was selectively upregulated during the convalescent phase (Fig 1I). Together, these findings demonstrated that CHIKV infection promotes robust STAT1-mediated inflammation. Monocytes appear to contribute to an antiviral and inflammatory state driven by the JAK-STAT signaling axis, likely maintained by sustained expression of multiple interferons across both acute and convalescent phases.

Taken together, these findings indicate that, in the peripheral blood of pediatric patients with acute CHIKD, CHIKV infection drives monocyte activation and subsequent differentiation into macrophages through the engagement of multiple PRR pathways, including TLRs, RLRs, and inflammasomes. This response is characterized by constrained NF-κB-dependent inflammation alongside robust multi-IFN-STAT1-dependent antiviral and inflammatory activity, highlighting a potential immunopathological role for monocytes, IFNs, and STAT1 signaling in acute inflammation and arthralgia observed in CHIKD.

### 3.2. CHIKV Infection During Monocyte-to-Macrophage Differentiation Promotes Viral Persistence and Extensive Cellular Remodeling

We previously reported that human monocytes are permissive to CHIKV infection *in vitro* and mount robust inflammatory and antiviral responses (23)(37). However, whether CHIKV infection directly impacts monocyte-to-macrophage differentiation remains to be elucidated. Building upon our *in vivo* characterization of monocyte responses in CHIKD patients (Fig 1), we differentiated primary human monocytes into macrophages either in the absence of virus (uninfected MDM) or after CHIKV challenge at an MOI of 10 (MDM-CHIKV). We then conducted comprehensive phenotypic, transcriptomic, and functional analyses, as outlined in Fig 2A.

**Figure 2.**
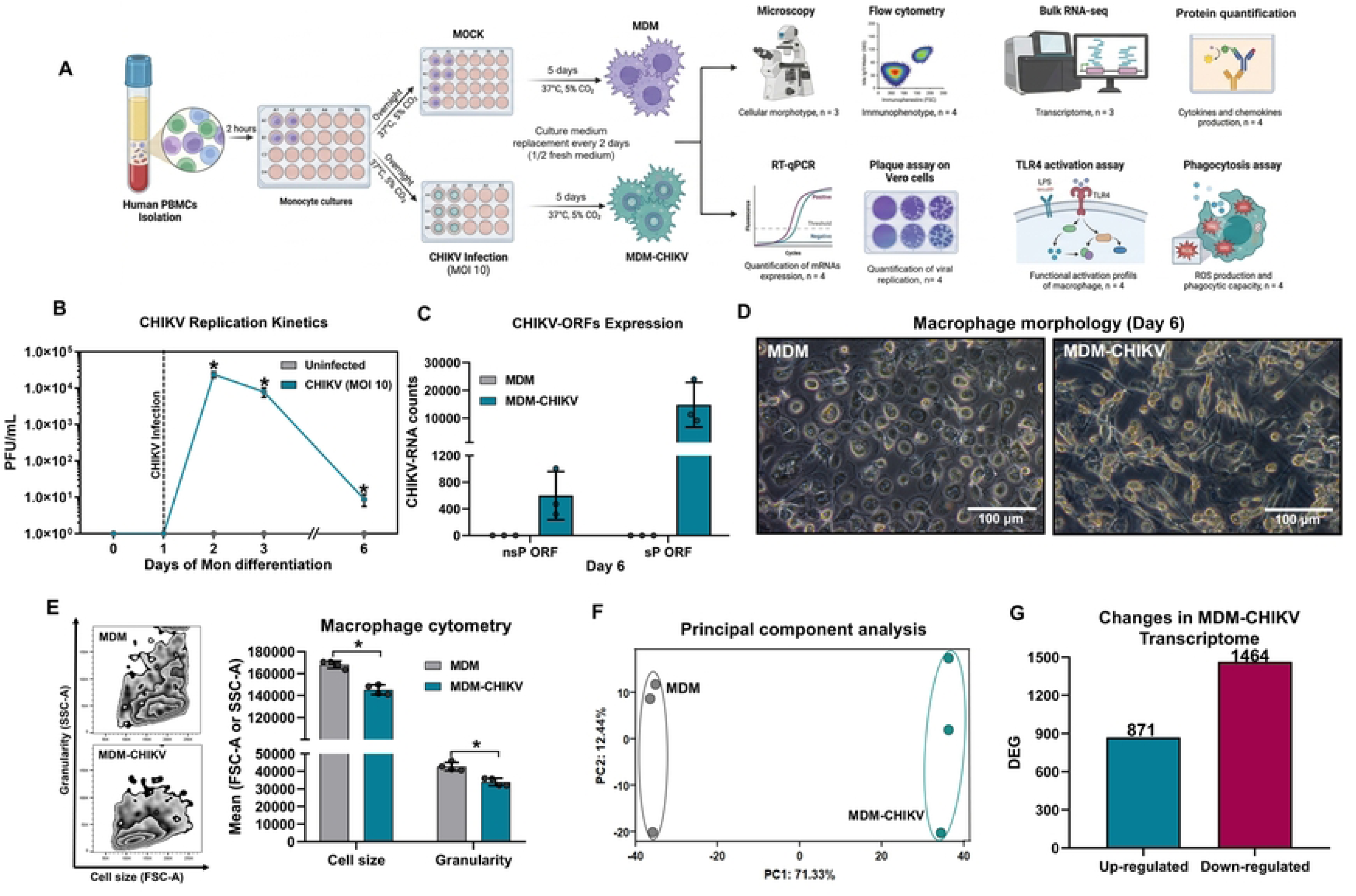
CHIKV infection during monocyte-to-macrophage differentiation promotes viral persistence and extensive cellular and transcriptomic remodeling. **A)** Schematic overview of the experimental workflow. **B)** Production of infectious CHIKV particles measured by plaque assay in Vero cells. Normality was assessed by the Shapiro-Wilk test, and sample comparisons were performed using the Mann-Whitney test, whereby * = p < 0.05 comparing uninfected and CHIKV-infected groups. **C)** Viral RNA quantification in infected MDM by bulk RNA-seq. **D)** Representative phase-contrast microscopy images showing distinct macrophage morphotypes. **E)** Representative flow cytometry plots and quantitative analysis of macrophage size and cytoplasmic granularity. Data are presented as mean ± SD. Normality was assessed using the Shapiro-Wilk test, and sample comparisons were performed by the Mann-Whitney test, whereby * = p < 0.05 comparing uninfected and CHIKV-infected groups. **F)** Principal component analysis of transcriptomic profiles. **G)** Summary of DEGs in MDM-CHIKV compared to uninfected control, identified using DESeq2 with a p-value < 0.05 and |log2 fold change (MDM-CHIKV vs. unstimulated MDM) | > 0.6.

We first confirmed the permissiveness of primary human monocytes to CHIKV replication. Consistent with our previous findings (23), viral replication was rapid, peaking at 1 dpi and declining significantly by day 2 (Fig 2B). Notably, by day 6, we still detected persistent release of infectious viral particles in the supernatants of MDM-CHIKV cultures (Fig 2B). Bulk RNA-seq analysis further confirmed the expression of ORFs encoding both non-structural (nsP) and structural (sP) viral proteins in MDM-CHIKV (Fig 2C). While nsP ORF transcript levels remained relatively low, sP ORF abundance was markedly higher. This pronounced shift toward sP ORF expression over nsP aligns with the canonical replication strategy of alphaviruses, demonstrating persistent infection and active viral RNA replication in macrophages differentiated from virus-challenged monocytes.

Morphological characterization at day 6 revealed that MDM-CHIKV cells acquired an elongated, rod-like, or ameboid morphology with prominent cytoplasmic projections compared to uninfected MDM (Fig 2D). Furthermore, MDM-CHIKV cells exhibited a significant reduction in cell size and cytoplasmic granularity (Fig 2E), indicating that CHIKV infection impairs normal MDM differentiation and induces distinct structural alterations. These morphological changes were accompanied by extensive transcriptional remodeling. PCA of bulk RNA-seq data showed clear separation between MDM and MDM-CHIKV transcriptomes, with PC1 accounting for 71.33% of the variance driven by infection status, and PC2 (12.44%) reflecting donor-to-donor variability (Fig 2F). Differential expression analysis identified 871 significantly upregulated and 1464 downregulated genes in MDM-CHIKV relative to uninfected MDM (Fig 2G). Collectively, these results demonstrate that CHIKV infection during monocyte-to-macrophage differentiation leads to persistent viral replication, altered cellular morphotypes, reduced cell size and internal complexity, and extensive transcriptomic reprogramming. These *in vitro* findings recapitulate key features observed in CHIKD patients (Fig 1) and support a model in which CHIKV skews monocyte differentiation toward a chronically infected, phenotypically altered macrophage state. However, whether this reprogramming reflects a classical (M1) or alternative (M2) polarization spectrum in MDM-CHIKV cells remains to be fully defined.

### 3.3. Persistent CHIKV Infection During Monocyte-to-Macrophage Differentiation Drives M1-Like Polarization, IFN-π/IL-27 Production, STAT1-Dependent Inflammation, and Enhanced Antigen Presentation

Changes in macrophage morphology and cellular features are frequently accompanied by functional transcriptomic reprogramming. We therefore hypothesized that CHIKV infection during monocyte-to-macrophage differentiation not only alters cellular morphology but also impacts the activation profile and functional capacity of MDM-CHIKV compared to uninfected MDM. Given our *in vivo* observation that CHIKD patients exhibit robust multi-IFN-STAT1-dependent inflammatory responses, we first evaluated the capacity of MDM-CHIKV cells to produce IFNs. Interestingly, we detected modest but significant induction of *IFNB1* mRNA, but not of *IFNA2*, *IFNE*, *IFNG*, or *IFNL1* (Fig 3A). In contrast, both subunits of the IFN-π/IL-27 heterodimer (*IL27p28* and *EBI3*) were markedly upregulated (Fig 3A). Furthermore, IFN-π/IL-27 protein secretion was significantly elevated in MDM-CHIKV culture supernatants (Fig 3B), suggesting that MDM-CHIKV cells may promote IFN-π/IL-27-dependent JAK-STAT activation. Consistent with this, MDM-CHIKV supernatants contained high levels of STAT1-responsive chemokines, including CCL-2 and CCL-5 (Fig 3C). Additionally, MDM-CHIKV cells produced elevated levels of NF-κB-dependent cytokines, including TNF-α and IL-6 (Fig 3D), indicating that persistent CHIKV infection induces a distinct, multifactorial inflammatory polarization state.

**Figure 3.**
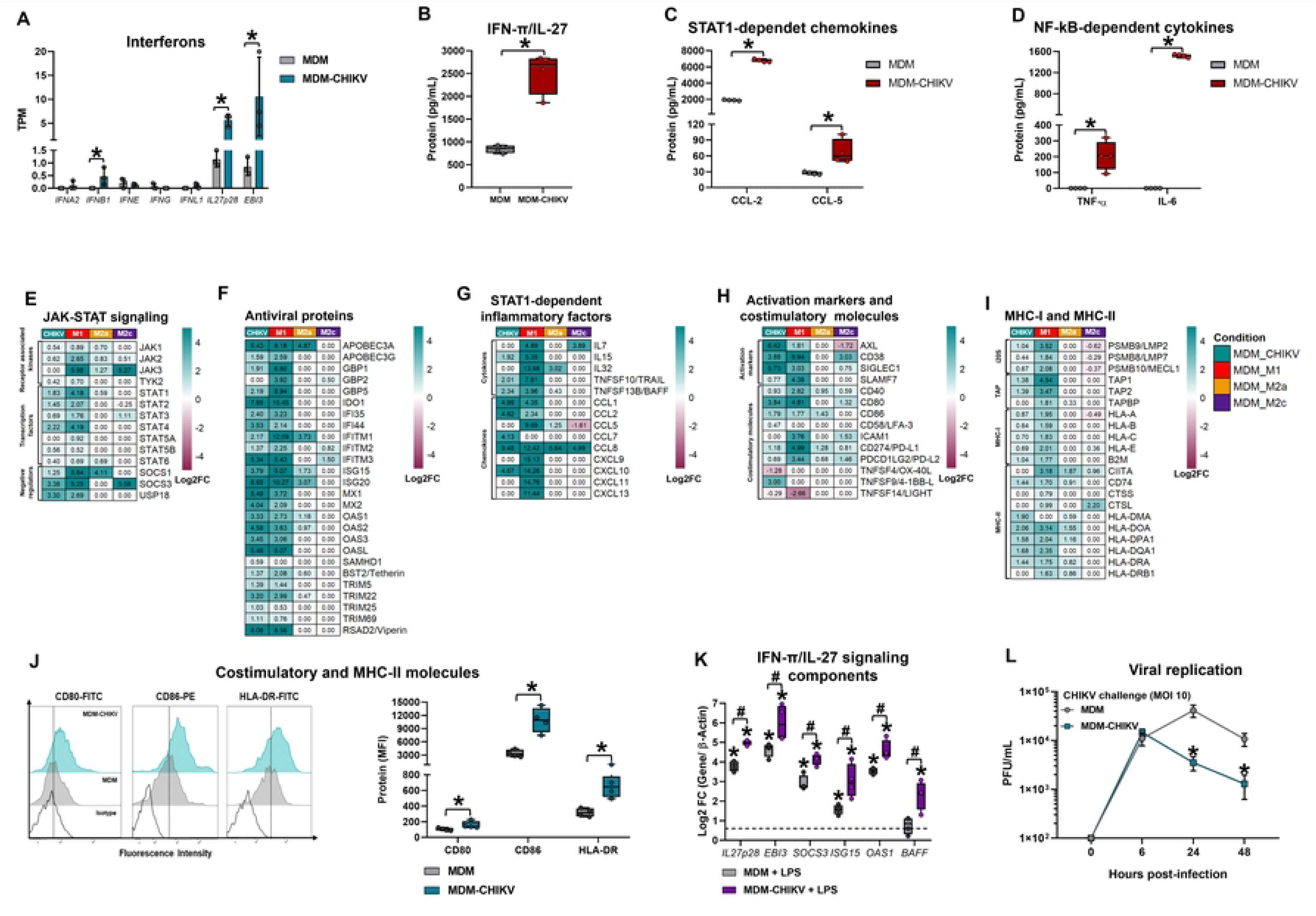
Persistent CHIKV infection during monocyte-to-macrophage differentiation drives M1-like polarization, IFN-π/IL-27 production, STAT1-dependent inflammation, and enhanced antigen presentation. **A)** Relative mRNA expression of human interferons in uninfected and MDM-CHIKV. **B-D)** Supernatant protein levels of **B)** IFN-π/IL-27, **C)** STAT1-dependent inflammatory factors, and **D)** NF-κB-dependent mediators evaluated by ELISA. Normality was assessed using the Shapiro-Wilk test, whereby * = p < 0.05 comparing uninfected and CHIKV-infected groups. **(E–I)** DEG were identified using DESeq2 with a p-value < 0.05 and |log2 fold change (MDM-CHIKV or reference polarized MDM vs. unstimulated MDM)|> 0.6. **E)** JAK-STAT signaling, **F)** antiviral ISGs, **G)** STAT1-dependent inflammatory factors, **H)** activation and costimulatory markers, and **I)** MHC-I and MHC-II machinery. **J)** Representative histograms and surface expression analysis of CD80, CD86, and HLA-DR by flow cytometry. Normality was assessed using the Shapiro-Wilk test, and sample comparisons were performed by the Mann-Whitney test, whereby * = p < 0.05 comparing uninfected and CHIKV-infected groups. **K)** RT-qPCR expression of IFN-γ/IL-27 signaling components under basal and LPS-stimulated conditions (24 h). Normality was assessed using the Shapiro-Wilk test, and sample comparisons were performed using the Kruskal-Wallis test with Dunn’s post-hoc test, whereby *= p < 0.05 comparing LPS-stimulated and Mock macrophages, and # = p < 0.05 comparing LPS-stimulated MDM with LPS-stimulated MDM-CHIKV. **L)** Infectious viral release in culture supernatants measured by plaque assay on Vero cells. Data represent medians with interquartile ranges, whereby * = p < 0.05 comparing uninfected/unstimulated control vs. LPS-stimulated MDMs using Mann–Whitney test or Kruskal–Wallis test with Dunn’s post-hoc correction.

To systematically define the polarization profile of MDM-CHIKV, we performed a comparative transcriptomic analysis using the publicly available bulk RNA-seq dataset (GSE205990) generated by Embregts et al. (43), in which human MDM were polarized for 48 hours into M1 (IFN-γ + LPS), M2a (IL-4), or M2c (IL-10) states. We first examined components of JAK-STAT signaling pathways, a key regulator of macrophage polarization. Both MDM-CHIKV and reference MDM-M1 cells exhibited robust transcriptional upregulation of key mediators of the JAK-STAT cascade, albeit with differing expression magnitudes. Specifically, significant induction was observed for *JAK2*, *STAT1*, *STAT2*, *STAT3*, and *STAT4*, alongside canonical negative regulators, including *SOCS1*, *SOCS3*, and *USP18* (Fig 3E). In contrast, MDM-M2a cells selectively upregulated *JAK1*, *JAK2*, *JAK3*, and *STAT6* with modest *STAT1* and high expression of *SOCS1*, whereas MDM-M2c cells preferentially expressed *JAK3*, *STAT3*, and *SOCS3* mRNA (Fig 3E). These findings validate lineage-appropriate JAK-STAT activation in reference polarized macrophages and position the MDM-CHIKV transcriptome closest to the M1 profile.

We also observed that MDM-CHIKV and MDM-M1 cells showed high transcriptional convergence, with both populations exhibiting robust upregulation of ISGs encoding canonical antiviral effectors, including *APOBEC3A/G*, *GBP1/2/5*, *IFI35/44*, *IFITM1-3*, *MX1/2*, *OAS1/2/3/L*, and *TRIM5/22/25/69*, alongside *IDO1*, *ISG15*, *ISG20*, *BST2/*Tetherin, and *RSAD2*/Viperin (Fig 3F). Regarding MDM-M2a, the number of ISGs expressed in these cells was low, with only six induced genes (*APOBEC3A*, *IFITM1*, *ISG15*, *ISG20*, *OAS1*, and *OAS2*), whereas MDM-M2c cells selectively expressed *IFITM2* and *IFITM3* (Fig 3F).

We next evaluated STAT1-dependent inflammatory mediators. Both MDM-CHIKV and MDM-M1 cells significantly upregulated transcript levels of *IL15*, *TNFSF10/TRAIL*, *TNFSF13B/BAFF*, *CCL1*, *CCL2*, *CCL8*, and *CXCL10* (Fig 3G), confirming shared activation of a STAT1-driven inflammatory program. However, MDM-M1 cells also induced *IL7*, *IL32*, *CCL5*, *CXCL9*, *CXCL11*, and *CXCL13* (Fig 3G), reflecting a broader pro-inflammatory spectrum than that of MDM-CHIKV. By comparison, MDM-M2a macrophages upregulated only *IL32*, *CCL5*, and *CCL8*, whereas MDM-M2c macrophages induced only *IL7* and *CCL8* (Fig 3G), validating the attenuated STAT1 response in M2-polarized states.

Beyond driving inflammation, STAT1 activation promotes antigen presentation capacity (15)(16). We therefore assessed expression of costimulatory molecules and components of the antigen-processing machinery. As shown in Figure 3H, MDM-CHIKV cells closely aligned with reference MDM-M1 macrophages, as both upregulated key immune activation markers, including AXL receptor tyrosine kinase (*AXL)*, *CD38*, Sialic acid binding Ig-like lectin 1 (*SIGLEC1)*, SLAM family member 7 (*SLAMF7),* alongside costimulatory and immune-checkpoint molecules such as *CD40*, *CD80*, *CD86*, *CD274/*PD-L1, and *PDCD1LG2*/PD-L2. Notably, MDM-M2a and MDM-M2c macrophages also expressed subsets of these factors, albeit at lower levels, including *CD40*, *CD274*, and *PDCD1LG2*, with MDM-M2c cells further inducing *CD38*, *SIGLEC1*, *CD80*, and *ICAM1* (Fig 3H). These results indicate that while MDM-CHIKV cells display a prominent M1-like costimulatory profile, the expression of individual costimulatory markers is not exclusively restricted to M1 polarization.

Regarding antigen processing, both MDM-CHIKV and MDM-M1exhibited robust upregulation of transcripts encoding key subunits of 20S immunoproteasome (i20S), including low molecular weight polypeptide 2 (*PSMB9/*LMP2), *PSMB8/*LMP7, and the multi-catalytic endopeptidase complex-like 1 (*PSMB10/*MECL1); TAP transporters such as *TAP1*, *TAP2*; and MHC-I molecules, including *HLA-A*, *HLA-B*, *HLA-C*, *HLA-E*, and Beta-2 microglobulin (*B2M*) (Fig 3I). Conversely, components of the MHC-II presentation machinery, including the master Class II major histocompatibility complex transactivator (*CIITA*), *CD74*, *HLA-DOA*, *HLA-DPA*, and *HLA-DRA*, were highly upregulated in MDM-M1 cells, whereas MDM-CHIKV and MDM-M2a displayed partial induction of this pathway (Fig 3I), suggesting broad potential for T-cell presentation across these subsets. In contrast, MDM-M2c cells failed to induce MHC-II components, downregulated i20S subunits and *HLA-A*, yet selectively upregulated *CIITA* and *CTSL* (Fig 3I). Validating these transcriptomic findings at the protein level, MDM-CHIKV cells exhibited significantly higher surface expression of CD80, CD86, and HLA-DR compared to uninfected MDM (Fig 3J), confirming a functional M1-like activation phenotype.

Having observed that MDM-CHIKV exhibits an M1-like and ISG-enriched transcriptional profile, we next asked whether persistent infection alter their capacity to respond to innate immune stimuli. To this end, we stimulated MDM and MDM-CHIKV cells with ultrapure LPS (100 ng/mL), a selective TLR4 agonist. Remarkably, MDM-CHIKV cells mounted a heightened antiviral response, further upregulating both subunits of IFN-π/IL-27 (*IL27p28* and *EBI3*), *SOCS3*, and downstream ISGs, including *ISG15*, *OAS1*, and *TNFSF13B*/BAFF (Fig 3K). These results indicate that MDM-CHIKV retains full capacity to mount robust IFN-π/IL-27-STAT1-dependent pro-inflammatory and antiviral responses upon TLR4 engagement.

Given the ISG-enriched antiviral state of MDM-CHIKV, we next asked whether persistent infection limits their susceptibility to reinfection. To test this, we subjected MDM-CHIKV to a secondary challenge with CHIKV (MOI 10), viral replication kinetics were comparable between MDM-CHIKV and primary MDM cells during the first 6 h.p.i. (Fig 3L). However, in MDM-CHIKV cells, viral titers reached a maximum peak at 6 h.p.i, after which viral replication declined sharply and continued to decrease through 48 h.p.i. In contrast, MDM cells exhibited a peak in viral replication at 24 h.p.i, followed by a gradual decline that remained above the low levels observed in MDM-CHIKV at later time points (Fig 3L). These kinetic differences suggest that MDM-CHIKV and MDM cells primarily diverge in their capacity to sustain viral replication over time, rather than in their initial permissiveness. This observation supports the notion that the IFN-π/IL-27-dependent antiviral program restricts secondary viral replication without eliminating viral persistence. Collectively, these results demonstrate that MDM-CHIKV cells adopted an M1-like polarization state characterized by IFN-π/IL-27 production, STAT1-dependent antiviral and inflammatory responses, and marked upregulation of antigen-processing and presentation machinery. This functional profile positions MDM-CHIKV cells as potent activators of T-cell responses, distinguishing them from classically M2-polarized macrophages and revealing a previously unrecognized functional consequence of CHIKV-driven monocyte differentiation.

### 3.4. MDM-CHIKV Exhibit Constitutive M1-Like PRR-NF-κB Activation but Adopt an M2-Associated Regulatory Profile Upon TLR4 Engagement

Macrophage polarization dynamically modulates PRR expression, thereby shaping pathogen-sensing capacity and downstream inflammatory output (27)(52). To further define the transcriptomic identity of MDM-CHIKV, we benchmarked its PRR signaling profile against classical polarization states (MDM-M1, MDM-M2a, and MDM-M2c). Transcriptomic profiling revealed distinct TLR expression patterns across subsets. While MDM-M1 cells significantly upregulated *TLR1*, *TLR2*, *TLR4*, and *TLR8*, with concurrent downregulation of *TLR5*, MDM-CHIKV cells specifically upregulated *TLR2*, *TLR3*, *TLR4*, and *TLR8*, receptors key for sensing viral nucleic acids and viral structural proteins (Fig 4A). Furthermore, both MDM-M1 and MDM-CHIKV displayed significant upregulation of critical adaptors and signaling mediators, including *CD14*, TIR-domain-containing adapter-finducing interferon-β (*TICAM1*/TRIF), IL1R-associated kinase 2 (*IRAK2)*, *IRAK3*, and *IRAK4* (Fig 4A), indicating a heightened intrinsic potential for TLR-dependent signal transduction. In contrast, MDM-M2a cells significantly downregulated *TLR1*, *CD14*, *MyD88*, *IRAK1*, and *IRAK3* (Fig 4A), consistent with IL-4-mediated suppression of TLR responsiveness. Notably, MDM-M2c cells exhibited elevated expression of *TLR1*, *TLR2*, *TLR7*, and *TLR8*, and *CD14* (Fig 4A), highlighting a distinct pathogen-sensing repertoire in this alternative regulatory phenotype.

**Figure 4.**
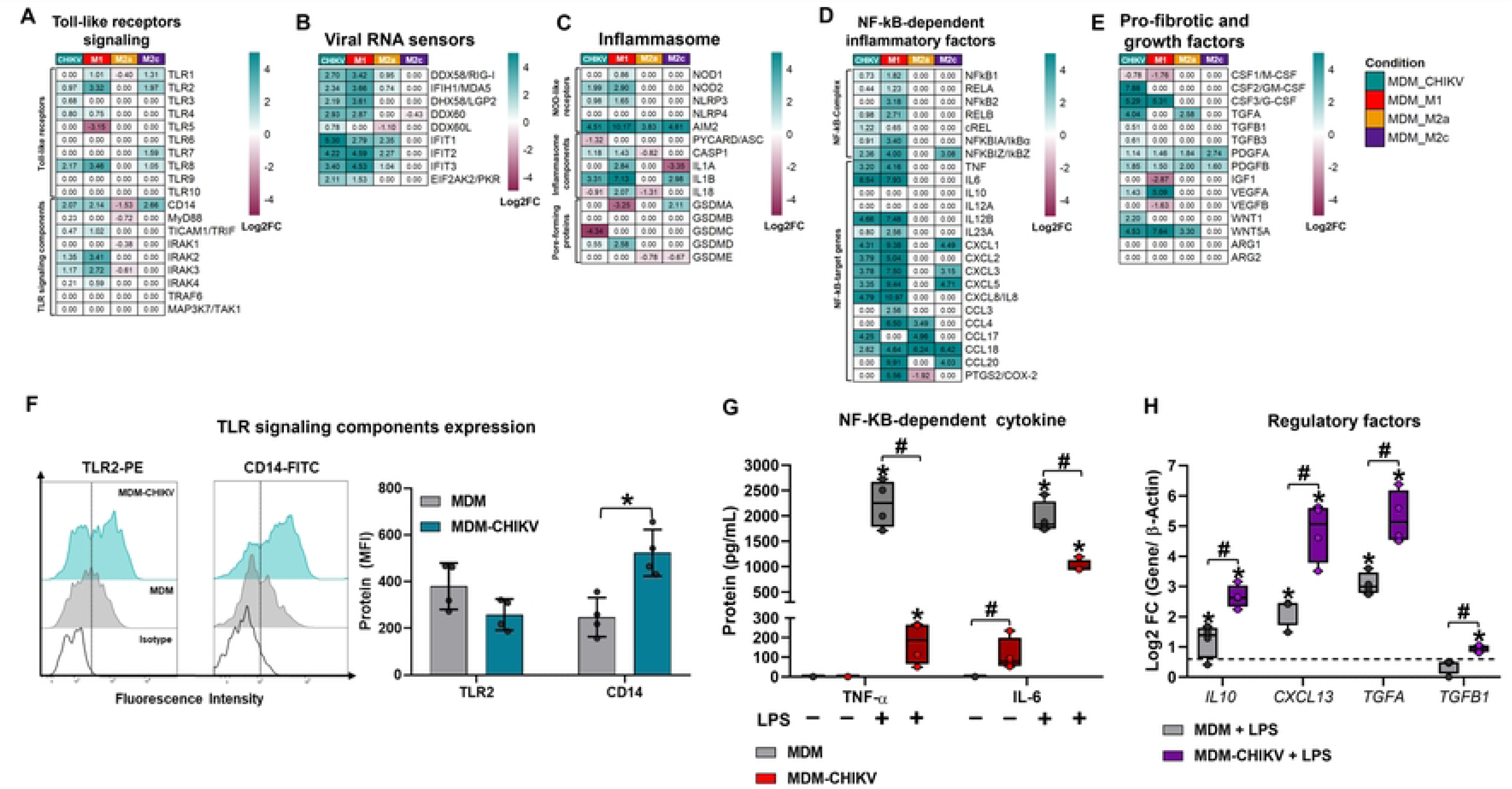
MDM-CHIKV cells exhibit constitutive M1-like PRR–NF-κB activation but adopt an M2-associated regulatory profile upon TLR4 engagement. Primary human monocyte cultures were either infected with CHIKV or not and differentiated into MDM as outlined in Fig. 2A. **(A-E)** DEGs were identified using DESeq2 with a p-value < 0.05 and |log2FC (MDM-CHIKV or reference polarized MDM subsets vs. unstimulated MDM)|> 0.6: **A)** TLR signaling components, **B)** Viral RNA sensors, **C)** Inflammasome elements, **D)** NF-κB-dependent inflammatory factors, and **E)** Pro-fibrotic and growth factors. **F)** Representative histograms and flow cytometric quantification of surface TLR2 and CD14 expression. Normality was assessed using the Shapiro-Wilk test, and sample comparisons were performed using the Mann-Whitney test, whereby * = p < 0.05 comparing MDM and MDM-CHIKV groups. **(G, H)** Unstimulated or LPS-stimulated (24 h) MDM and MDM-CHIKV cultures were evaluated for: **G)** Protein levels of NF-κB-dependent cytokines by ELISA, and **H)** mRNA expression of regulatory/anti-inflammatory factors by RT-qPCR. Normality was assessed using the Shapiro-Wilk test, and sample comparisons were performed using the Kruskal-Wallis test with Dunn’s post-hoc test, whereby *= p < 0.05 comparing LPS-stimulated and Mock macrophages, and # = p < 0.05 comparing LPS-stimulated MDM with LPS-stimulated MDM-CHIKV.

Consistent with these findings, macrophage polarization differentially modulated RIG-like receptor expression. Both MDM-CHIKV and MDM-M1 cells exhibited significantly elevated transcript levels of core viral RNA sensors, including *DDX58*/RIG-I, *IFIH1*/MDA5, *DHX58/*LGP2, *DDX60*, *IFIT1-3*, and *EIF2AK2*/PKR (Fig 4B). MDM-M2a cells also upregulated a subset of these RNA sensors (*DDX58*, *IFIH1*, and *IFIT1-3* (Fig 4B), suggesting that IL-4 stimulation partially enhances intrinsic antiviral potential. In contrast, MDM-M2c cells showed no significant induction of RLRs; instead, they showed a downregulation of *DDX60* mRNA (Fig 4B). Beyond RLRs, evaluation of NOD-like receptors and inflammasome components revealed that both MDM-CHIKV and MDM-M1 cells markedly induced *NOD2*, *NLRP3*, and *AIM2*, together with core inflammasome effectors, including *CASP1* and *IL1B* (Fig. 4C), pointing toward active inflammasome priming in both macrophage subsets (Fig 4C). However, MDM-M1 cells additionally upregulated *IL1A*, *IL18*, and *GSDMD* (Fig 4C), defining a fully equipped inflammasome profile with high potential to drive pyroptotic cell death. In contrast, MDM-M2a cells induced *AIM2* but significantly downregulated *CASP1*, *IL18*, and *GSDME* (Fig 4C). Similarly, MDM-M2c upregulated *AIM2*, *IL1B*, and *GSDMA* while downregulating *IL1A* and *GSDME* (Fig 4C), indicating subset-specific modulation of inflammasome execution pathways.

In line with significant PRR induction, both MDM-CHIKV and MDM-M1 cells markedly upregulated key NF-κB-complex components, including *NFKB1*, *RELA*, *RELB*, c*REL*, *NFKBIA/*IkBα, and *NFKBIZ*/IkBZ (Fig 4D). This transcriptional activation was accompanied by robust induction of downstream NF-κB-dependent inflammatory mediators, including *TNF*, *IL6*, IL-23 subunits (*IL12B* and *IL23A*), *CXCL1*-*3*, *CXCL5, CXCL8*/IL-8, and *CCL18* (Fig 4D). Notably, neither *IL10* nor *IL12A* showed induction, thereby confirming a sustained, baseline NF-κB-driven pro-inflammatory state. However, MDM-M1 also upregulated *NFKB2*, *CCL3*, *CCL4*, *CCL20,* and cyclooxygenase 2 (*PTGS2*/COX-2) (Fig 4D), indicating a stronger inflammatory response than that of MDM-CHIKV cells. MDM-M2a cells only upregulated *CCL4, CCL4, CCL18* and downregulated *PTGS2*. Particularly, *CCL17* mRNA was induced by both MDM-CHIKV and MDM-M2a cells, whereas MDM-M2c cells induced *NFKBIZ/*IkBZ, *CXCL1*, *CXCL3*, *CXCL5, CCL18*, and *CCL20* (Fig 4D), indicating that M2 phenotypes sustain a more restricted, chemokine-biased inflammatory output.

Furthermore, MDM-CHIKV and MDM-M1 cells exhibited a similar pattern of growth factor modulation, characterized by downregulation of *CSF1*/M*-CSF* and upregulation of granulocyte colony-stimulating factor (*CSF3*/G-CSF), platelet-derived growth factor subunit A and B (*PDGFA*/*B*), Vascular endothelial growth factor A (*VEGFA*), and Wnt family member 5A (*WNT5A*) (Fig 4E). Notably, MDM-CHIKV cells also upregulated a distinct set of immunomodulatory and pro-angiogenic factors, including *CSF2*/GM*-*CSF, transforming growth factor alpha (*TGFA*), *TGFB1*, *TGFB3*, and *WNT1* (Fig 4E). In contrast, MDM-M1 cells selectively down-regulated *IGF1* and *VEGFB*, suggesting that CHIKV-exposed macrophages acquire a specialized transcriptional profile that bridges pro-inflammatory activation with tissue-remodeling and regulatory responses compared to classical MDM-M1 macrophages. Meanwhile, MDM-M2 subsets display a more restricted growth factor profile. MDM-M2a cells induce *TGFA*, *PDGFA/B*, and *WNT5A*, whereas MDM-M2c cells selectively upregulate *PDGFA* and *PDGFB*, growth factors expressed across all four macrophage phenotypes evaluated (Fig 4E). Notably, none of the MDM polarization states induced the expression of the pro-fibrotic markers *ARG1* and *ARG2* (Fig 4E).

Consistent with our transcriptomic data, flow cytometric analysis confirmed that MDM-CHIKV cells exhibited significantly higher surface CD14 expression than uninfected MDM, alongside reduced TLR2 levels (Fig 4F), indicating active viral tuning of TLR pathway components. While MDM showed no baseline secretion of TNF-α or IL-6, MDM-CHIKV cells exhibited elevated IL-6 production (Fig 4G). Strikingly, upon LPS stimulation, MDM-CHIKV cells produced fewer NF-κB-dependent pro-inflammatory cytokines, specifically TNF-α and IL-6, than MDM (Fig 4G). This demonstrates that while MDM retain a robust capacity for classical TLR4-driven activation, CHIKV infection reprograms macrophages into a distinct activation state that is less responsive to secondary inflammatory challenges. Intriguingly, following LPS stimulation, MDM-CHIKV cells selectively upregulated mRNA expression of classic M2-associated regulatory and remodeling factors, including *IL10*, *CXCL13*, *TGFA*, and *TGFB1* (Fig 4H), indicating a secondary shift toward an M2-like immunosuppressive and tissue-protective phenotype. These findings further imply that full induction of M2-associated regulatory programs in alternatively activated macrophages may require functional PRR engagement to drive NF-κB-dependent transcriptional machinery, as observed in MDM-CHIKV. Taken together, these data reveal that CHIKV-exposed macrophages acquire a unique functional identity. While sharing core effector traits with classical M1 macrophages, such as broad PRR induction, baseline NF-κB activation, and robust pro-inflammatory cytokine and chemokine output, MDM-CHIKV cells distinctly retain *TLR3* expression, harbor an expanded growth factor repertoire, and undergo an M2-like regulatory shift upon TLR4 engagement, reflecting a hybrid, virus-adapted polarization state.

### 3.5. Complement System Activation and Phagocytic Receptor Modulation Define the Effector Profile of MDM-CHIKV, Shaping CHIKV-Associated Immunopathology

Phagocytosis is a primary effector function of macrophages; impairments in this process compromise pathogen clearance and the resolution of cellular debris, thereby fueling persistent inflammation (29,44). Because macrophage polarization critically determines phagocytic capacity and microbicidal activity, we analyzed the transcriptional landscape of key phagocytic receptors in MDM-CHIKV cells compared with classical (MDM-M1) and alternatively activated (MDM-M2a and MDM-M2c) macrophage subsets. Transcriptomic analysis revealed distinct differential regulation of C-type lectin receptors across macrophage subsets (Fig 5A). The macrophage mannose receptor (*MRC1*/CD206) was significantly upregulated in MDM-CHIKV, MDM-M2a, and MDM-M2c, whereas it was downregulated in MDM-M1 cells. While all evaluated macrophage populations displayed increased mRNA expression of dendritic cell-specific ICAM-3-grabbing non-integrin (*CD209*/DC*-*SIGN), only MDM-CHIKV cells uniquely upregulated the dendritic cell immunoreceptor (*CLEC4A*/DCIR). Conversely, whereas MDM-M1 cells selectively induced CLEC6A/*Dectin-2* and macrophage-inducible C-type lectin (*CLEC4E*/MINCLE), MDM-M2a cells downregulated these sensors along with *CLEC5A* (Fig 5A), highlighting divergent C-type lectin expression profiles between classical M1 and M2 subsets and the virus-adapted MDM-CHIKV phenotype.

**Figure 5.**
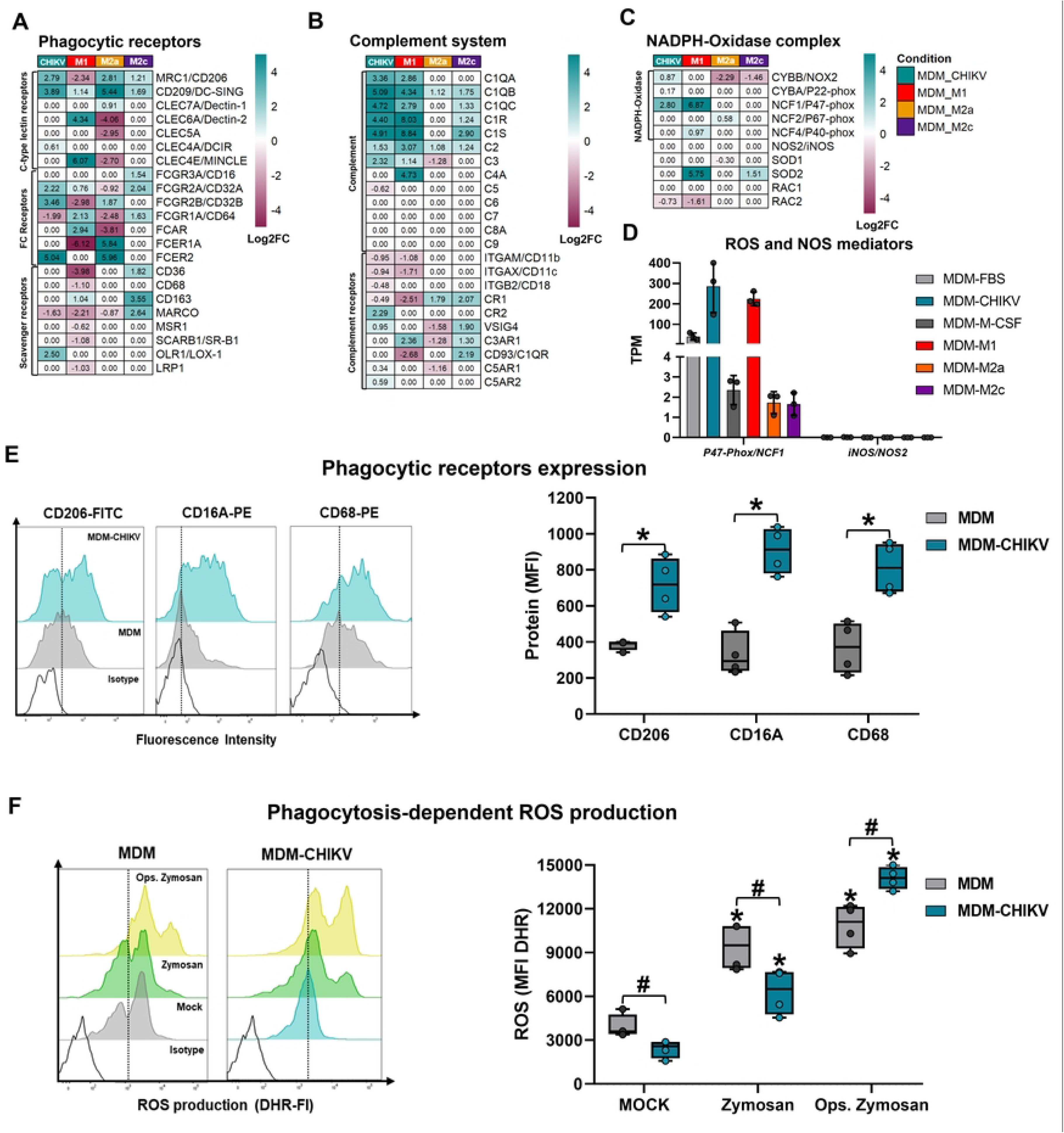
Complement system activation and phagocytic receptor modulation define the effector profile of MDM-CHIKV. Primary human monocyte cultures were either infected with CHIKV or left uninfected and then differentiated into MDMs as outlined in Fig. 2A. (**A-D**) DEGs were identified using DESeq2 with a p-value < 0.05 and |log2FC heatmaps compare mRNA expression profiles of MDM-CHIKV or reference polarized MDM subsets vs. unstimulated MDM)|> 0.6: A) Phagocytic receptors, **B)** Complement system components, and **C)** NADPH oxidase complex subunits. **D)** Relative mRNA expression (TPM) of ROS and NOS mediators. **E)** Representative histograms and surface expression analysis of CD206, CD16, and CD68 by flow cytometry. Data normality was assessed using the Shapiro-Wilk test. Differences between MDM and MDM-CHIKV groups were evaluated using the two-tailed Mann-Whitney test. Statistical significance was defined as * = p < 0.05. **F)** Flow cytometry quantification of intracellular ROS production in response to zymosan or opsonized zymosan stimulation (n = 4). Data represent medians with interquartile ranges. Statistical significance among unstimulated, zymosan-stimulated, and opsonized zymosan-stimulated macrophages is indicated by * = p < 0.05, whereas statistical significance between MDM and MDM-CHIKV groups is indicated by # = p < 0.05).

Expression of Fc receptors (FcRs) also varied markedly across polarization states (Fig 5A). Both MDM-CHIKV and MDM-M2a significantly upregulated the inhibitory Fcγ receptor *FCGR2B*/CD32B and *FCER2*, whereas MDM-M1 downregulated *FCGR2B*/CD32B and *FCER1A* (Fig 5A). MDM-M1 additionally upregulated the high-affinity IgG receptor *FCGR1A*/CD64 and the IgA receptor *FCAR*, both of which were downregulated in MDM-M2a cells but not in MDM-M2c (Fig 5A). Furthermore, *FCGR2A*/CD32A was co-upregulated in MDM-CHIKV and MDM-M2c cells.

Scavenger receptor expression was also differentially regulated across subsets. MDM-M1 strongly repressed an array of scavenger receptors, including *CD36*, *CD68*, *MARCO*, *MSR1*, Scavenger receptor class B member 1 (*SCARB1*/SR*-B1*), and LDL receptor-related protein 1 (*LRP1*) (Fig 5A), consistent with a specialized microbicidal rather than homeostatic/tissue-remodeling phenotype. In contrast, MDM-M2c cells upregulated *CD36*, *CD163*, and *MARCO* (Fig 5A), reflecting a classical reparative profile. Notably, MDM-CHIKV cells selectively upregulated oxidized low-density lipoprotein receptor 1 (*OLR1*/LOX-1) (Fig 5A), whereas MDM-M2a showed no significant induction of scavenger receptors and downregulated *MARCO* mRNA. Collectively, these findings indicate that phagocytic and scavenger receptor expression in MDM-CHIKV cells is highly context-dependent, blending M2-associated FcR profiles with a distinct, LOX-1-driven scavenger signature.

The complement system is a pivotal innate immune cascade that promotes phagocytosis and pathogen clearance; however, its persistent or aberrant activation is tightly associated with tissue injury and chronic immunopathology (53). To determine how CHIKV infection and macrophage polarization shape this system, we profiled the transcriptional expression of complement cascade components across macrophage subsets (Fig 5B). Both MDM-CHIKV and MDM-M1 cells shared a robust complement-activation signature characterized by marked upregulation of the C1q subunits (*C1QA*, *C1QB*, *C1QC*), initiation factors (*C1R*, *C1S*, and *C2*), and the central component *C3* (Fig 5B). Furthermore, MDM-M1 cells uniquely expressed *C4A*, indicating concurrent engagement of both the classical/lectin and alternative pathways. While MDM-M2c cells also upregulated several of these transcripts, MDM-M2a cells displayed a more restricted complement profile, characterized by upregulation of *C1QB* and *C2* alongside the downregulation of *C3* (Fig 5B). Together, these data demonstrate that MDM-CHIKV cells acquire a specialized, pro-inflammatory complement expression pattern that closely mimics the M1 transcriptional program, whereas M2a subsets exhibit a limited intrinsic capacity for complement cascade amplification. Strikingly, none of the macrophage subtypes modulated the expression of terminal membrane attack complex (MAC) components, including *C5*, *C6*, *C7*, *C8A*, and *C9* (Fig. 5B), indicating that human macrophages alone are insufficient to support terminal lytic complement activation.

Regarding complement receptors involved in phagocytosis, both MDM-CHIKV and MDM-M1 cells significantly downregulated complement receptor 3 (CR3) *alpha* integrin (*ITGAM*/CD11b), CR4 *alpha*-integrin (*ITGAX*/CD11c), and complement receptor 1 (*CR1*) (Fig. 5B). Intriguingly, MDM-CHIKV were the only macrophages that upregulated complement C3d receptor 2 (*CR2*/CD21) and downregulated *ITGB2*/CD18 (Fig. 5B), whereas MDM-M1 selectively induced *C3AR1* and suppressed the C1q receptor *CD93* (Fig. 5B). These findings suggested a distinct remodeling of complement-sensing machinery in CHIKV-exposed macrophages. Among M2 macrophages, both MDM-M2a and MDM-M2c cells significantly upregulated *CR1* (Fig. 5B). Notably, MDM-M2c exhibited markedly higher transcript levels of V-set and immunoglobulin domain containing 4 (*VSIG4*), *C3AR1*, and *CD93*(Fig. 5B), whereas MDM-M2a downregulated *VSIG4*, *C3AR1*, and *C5AR1* (Fig. 5B), underscoring a distinct complement-handling program within M2 sub-phenotypes. Together, these results demonstrate that MDM-CHIKV cells closely align with MDM-M1 cells in driving complement system activation and remodeling phagocytic/complement sensing machinery.

Consistent with modulation of phagocytic receptors and complement activation, we next evaluated the expression of NADPH oxidase complex components and reactive species-generating machinery. Both MDM-CHIKV and MDM-M1 cells significantly upregulated *NCF1/*p47-phox (Fig. 5C and D), while downregulating the small GTPase 2 (*RAC2*). Notably, only MDM-CHIKV significantly upregulated the catalytic core component cytochrome B-245 beta chain (*CYBB*/NOX2), whereas MDM-M2a and MDM-M2c cells downregulated its expression (Fig. 5C, D), indicating an enhanced intrinsic capacity for ROS production upon phagocytosis in CHIKV-exposed macrophages compared with the M2 phenotype. Additionally, both MDM-M1 and MDM-M2c cells upregulated superoxide dismutase 2 (*SOD2*), and MDM-M1 uniquely induced *NCF4* (p40-phox) (Fig. 5C), suggesting that MDM-M1 cells also maintain a robust ROS-generating capacity. Notably, none of the MDM populations displayed significant induction of nitric oxide synthase 2 (*NOS2*/iNOS) (Fig. 5C and D), in sharp contrast to the high constitutive expression of *NCF1* (Fig. 5D). This absence highlights a fundamental functional divergence between human and murine macrophages, as murine M1-macrophages classically rely on high NOS2 expression to drive reactive nitrogen species (RNS) production (54)(55). Taken together, these results demonstrate that MDM-CHIKV cells acquire a distinct phagocytic and microbicidal identity that integrates functional features of both M1 and M2 polarization states. This specialized profile, defined by the selective co-induction of specific phagocytic and scavenger receptors, complement cascade components, and NADPH oxidase subunits, positions CHIKV-exposed macrophages as a key dual-function effector capable of driving pathogen surveillance while simultaneously perpetuating inflammation-associated immunopathology.

### 3.6. MDM-CHIKV Cells Exhibit Enhanced Phagocytic Capacity and Robust ROS Production Upon the Phagocytosis of Immune Complexes

Based on our transcriptomic findings, we evaluated the surface phenotype and phagocytic potential of uninfected MDM and MDM-CHIKV cells. Flow cytometry analysis revealed that MDM-CHIKV cells significantly upregulated surface expression of CD206, CD16A/FCγRIIIA, and CD68 compared with uninfected MDM (Fig. 5E). This protein-level validation confirms that MDM-CHIKV cells adopt an M2-like immunophenotype enriched for key endocytic receptors and FcRs, indicating enhanced phagocytic capacity.

To functionally assess phagocytosis-associated effector responses, we used zymosan, a *Saccharomyces cerevisiae* cell wall preparation widely employed to model phagocytosis and respiratory burst activation in innate immune cells (29,44,45). Unopsonized zymosan is recognized by multiple PRRs, including TLR2/TLR6 heterodimers and C-type lectin receptors such as CD206, DC-SIGN, and Dectin-1 (29,44,45). Serum-opsonized zymosan (ZOP) engages additional opsonin-dependent pathways including Fcγ receptors and complement receptors, thereby accelerating phagocytic uptake, driving NADPH oxidase activation, and promoting robust superoxide anion (O₂⁻) generation (44). To functionally validate these transcriptional and surface receptor changes, we measured ROS production in MDM and MDM-CHIKV cells after challenge with either unopsonized zymosan or ZOP. Under basal conditions, unstimulated MDM-CHIKV cells (MOCK) displayed significantly lower baseline ROS levels than MOCK-untreated MDM (Fig. 5F). Upon zymosan stimulation, both macrophage populations produced significantly higher ROS levels relative to their respective MOCK-treated controls; however, zymosan-stimulated MDM mounted a more robust oxidative burst than MDM-CHIKV cells (Fig. 5F), indicating that unpolarized MDM mounts a more robust oxidative response to non-opsonized zymosan.

Notably, this functional hierarchy was completely reversed following stimulation with ZOP (ops. Zymosan). Under these conditions, MDM-CHIKV cells generated significantly higher ROS levels than uninfected control MDM (Fig. 5F), demonstrating that CHIKV infection primes macrophages for an enhanced oxidative burst specifically in response to opsonized targets and immune complexes. This functional divergence directly aligns with the differential upregulation of Fcγ and complement receptors identified in our transcriptomic and flow cytometry analyses. In summary, MDM-CHIKV cells acquired an M2-like immunophenotype marked by increased surface density of CD206, CD16, and CD68. Functionally, these cells display a specialized, context-dependent oxidative program: while attenuated in response to non-opsonized targets, they mount an amplified respiratory burst upon uptake of opsonized particles or immune complexes. This heightened reactivity to opsonized targets is fully consistent with their FCγ and complement receptor profile, positioning CHIKV-exposed macrophages for efficient clearance of opsonized entities while potentially contributing to inflammation-mediated tissue damage during CHIKV infection.

### 3.7. Single-Cell Transcriptomics Reveals Macrophages as the Primary Replicative Reservoir and Drivers of Stromal Remodeling in Chronic CHIKV-Arthritis

To further corroborate the biological relevance of MDM-CHIKV in the *in vivo* immunopathogenesis of CHIKV-induced arthritis, we reanalyzed the scRNA-seq dataset GSE300299 (33). This dataset comprises joint-associated tissues harvested from the ipsilateral ankle of immunocompetent wild-type (WT) C57BL/6J mice. In this well-established model, mice mount functional IFN-dependent antiviral responses yet develop chronic arthritis following CHIKV infection (33)(56).

We first characterized the transcriptional landscape within the arthritic joints (Fig 6A, B). In WT C57BL/6J mice, we observed a high abundance of joint stroma-associated cell populations, including synoviocytes (*Prg4^+^*, *Col1a1^+^*, *Col6a1^+^*), endothelial cells (*Pecam1^+^*, *Cdh5^+^*, *Eng^+^*), and skeletal muscle cells (*Myod1^+^*, *Myh2^+^*, *Dmd^+^*) (Fig. 6A, B). Additionally, we detected the infiltration of distinct immune cell populations, comprising myeloid lineage-specific subsets such as monocytes (*Ly6c2^+^*, *Ccr2^+^*, *Cd14^+^*), macrophages (*Adgre1^+^*/F4/80, *CD68^+^*, *CD163^+^*), and polymorphonuclear neutrophils (*Ly6g^+^*, *Itgam^+^*/Cd11b, *Cxcr2^+^*, *Lif^+^*) (Fig. 6A, B), together with lymphoid lineage-specific populations including NK cells (*Ncr1^+^*/Cd56, *Klrb1c^+^*, *Itga2^+^*), T cells (*Cd3d^+^*, *Cd4^+^*, *Cd8a^+^*), and B cells (*Cd19^+^*, *Cd79a^+^*, *Ms4a1^+^*) (Fig 6A, B).

**Figure 6.**
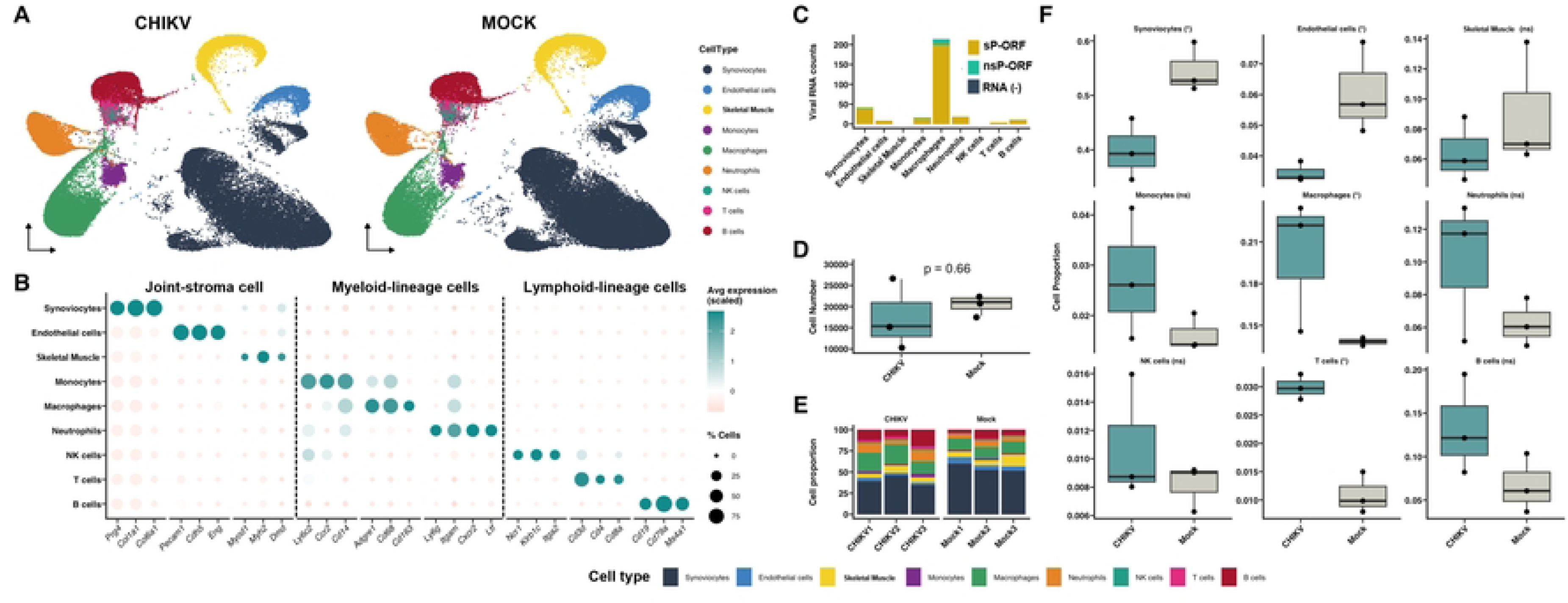
Single-Cell Transcriptomics Identifies Macrophages as the Primary Viral Reservoir and Drivers of Cellular Remodeling in Chronic CHIKV-induced Arthritis. **A)** UMAP projections of joint-associated cells from mock and chronically CHIKV-infected mice, colored by annotated cell type. **B)** Dot plot showing scaled average expression and percentage of expressing cells for canonical marker genes defining stromal, myeloid, and lymphoid populations. **C)** Viral RNA reads per cell type mapped to the structural polyprotein ORF (sP-ORF), nonstructural polyprotein ORF (nsP-ORF), and negative-strand viral RNA (-). **D)** Total single-cell yields per experimental condition. **E)** Joint cell-type composition across individual replicates. **F)** Relative proportion changes between mock and CHIKV-infected joints. Data are presented as medians with IQR (n = 3). Differences in cell type proportions between conditions were evaluated using the propeller function package in R. Proportions were logit- or arcsine square-root transformed before applying linear models and moderated t-tests. Statistical significance between CHIKV-infected mice and mock samples was set at FDR < 0.1 are indicated by * = p < 0.1; ns = not significant. n = 3.

We further identified that transcripts encoding the CHIKV subgenomic structural polyprotein (sP-ORF) were broadly detected across nearly all joint-associated cell populations in chronically infected mice, sparing only skeletal muscle cells and NK cells. Macrophages harbored the highest transcript abundance, followed by synoviocytes, neutrophils, and monocytes (Fig. 6C). In contrast, transcripts corresponding to the non-structural polyprotein (nsP-ORF), indicative of active viral replication, were strictly restricted to macrophages, with minor levels detected in synoviocytes and monocytes (Fig 6C). Furthermore, negative-sense viral RNA [RNA (−)], an essential intermediate of active viral replication, was uniquely detected within the macrophage population (Fig 6C). Together, these findings confirm that macrophages serve as the primary cellular reservoir for CHIKV within the joint microenvironment and represent the principal cell type supporting active viral replication during chronic infection.

In chronically CHIKV-infected WT C57BL/6J mice, total joint cell numbers remained comparable between mock- and CHIKV-infected animals at 28 dpi (Fig 6D). However, chronic CHIKV infection induced marked stromal remodeling, characterized by a significant reduction in the proportions of synoviocytes and endothelial cells (Fig 6E, F), a phenotype indicative of tissue structural deterioration and joint dysfunction. Concurrently, infected joints exhibited robust immune cell expansion and infiltration, marked by increased abundances of macrophages and T cells (Fig 6E, F). These cellular shifts reveal a transition toward a chronically inflamed microenvironment in which immune persistence is coupled with stromal degradation, ultimately driving long-term joint pathology.

### 3.8. Functional Profiling of Joint-Associated Macrophage Clusters Reveals Distinct Polarization States, Antigen-Presentation Capacity, and Inflammatory Programs

Subclustering of the joint-associated macrophage population resolved five transcriptionally distinct subsets (Fig 7A): Cluster 1 (47.4% in CHIKV vs. 13.0% in Mock), Cluster 2 (24.2% in CHIKV vs. 38.0% in Mock), Cluster 3 (19.7% in CHIKV vs. 21.8% in Mock), Cluster 4 (7.9% in CHIKV vs. 25.7% in Mock), and an Osteoclast cluster (0.7% in CHIKV vs. 1.4% in Mock). Notably, while transcripts corresponding to both nsP and sP viral ORFs were present in Cluster 1 and Cluster 2 macrophages, the RNA (−) replicative intermediate was exclusively restricted to Cluster 2 (Fig 7B). These findings identified Cluster-2 macrophages as the bona fide replicative viral reservoir within the joint’s microenvironment of chronically infected WT C57BL/6J mice.

**Figure 7.**
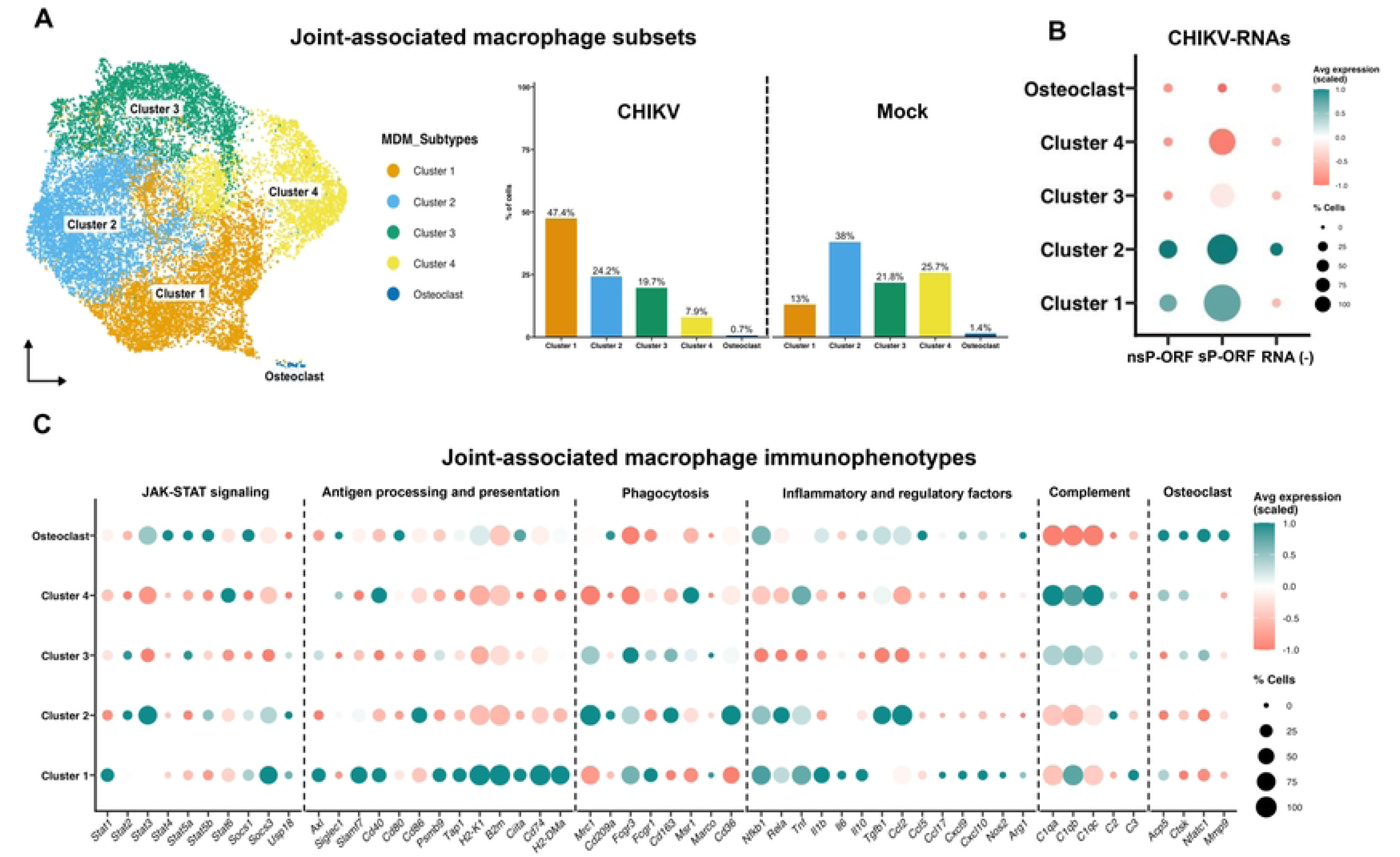
Functional Profiling of Joint-Associated Macrophage Clusters Reveals Distinct Polarization States, Antigen-Presentation Capacity, and Inflammatory Programs. **A)** UMAP projection of subclustered joint-associated macrophages (Cluster 1-4 and osteoclast) and their relative cell abundance (% of cells) in mock and CHIKV-infected joints. **B)** Dot plot showing viral RNA species (nsP-ORF, sP-ORF, and negative-sense RNA detected across macrophage clusters. **C)** Functional immunophenotyping of macrophage clusters illustrating scaled expression and prevalence of key gene sets, including JAK-STAT signaling, antigen presentation, phagocytosis, inflammation mediators, complement system components, and osteoclast-associated markers. In (B, C), dot color reflects scaled average expression and dot size indicates the percentage of expressing cells.

To define the functional profile and immunophenotype of the distinct joint-associated macrophage clusters, we evaluated a biomarker panel linked to phenotypic polarization and activation state. Cluster-1 exhibited high induction of *Stat1* alongside negative regulators of the IFN-dependent JAK-STAT signaling pathway, including *Socs1*, *Socs3*, and *Usp18* (Fig 7C). Consistent with a classical pro-inflammatory (M1) activation state this subset displayed elevated expression of costimulatory molecules (*Axl*, *Slamf7, Cd40*) and core components of the MHC class I and II peptide processing and presentation machinery (*Psmb9*, *Tap1*, *H2-K1*, *B2m*, *Ciita*, *Cd74*, and *H2-DMa*), suggesting a strong capacity for T-cells priming (Fig 7C). Furthermore, Cluster-1 macrophages displayed high expression of phagocytic receptors (*Fcgr3*/Cd16, *Fcgr1*/Cd64, and *Marco*), NF-κB-dependent inflammatory mediators (*Nfkb1*, *Rela*, *Tnf*, *Il1b*, *Il6*, *Il10*, *Ccl17*), STAT1-dependent chemokines/enzymes (*Cxcl9*, *Cxcl10*, and *Nos2/*iNOS), and complement components (*C1qb* and *C3*). This profile confirms that Cluster-1 macrophages adopt a classical M1-like pro-inflammatory phenotype, closely mirroring human M1 MDM (Figs 3–5).

In contrast, Cluster-2 macrophages upregulated *Stat2*, *Stat3*, and the regulators of the IFN-dependent JAK-STAT pathway, including *Socs1*, *Socs3*, and *Usp18* (Fig 7C). Unlike Cluster-1, these cells displayed minimal induction of costimulatory molecules, except for high *Cd86* expression, and low expression of antigen-presentation components, indicating limited T-cell activation potential. Strikingly, Cluster-2 macrophages coexpressed high levels of classical M2-assocaited phagocytic receptors (*Mrc1*/Cd206, *Cd209a*/Dc-sign, *Cd163*, and *Cd36*) alongside NF-κB-dependent factors (*Nfkb1*, *Rela*, and *Tnf*), *Tgfb1*, *Ccl2*, and *C2* (Fig 7C). This expression pattern reveals a hybrid M1/M2 activation state similar to that observed in human MDM-CHIKV cells (Figs 3–5).

Cluster-3 macrophages demonstrated elevated expression of *Stat2*, *Stat5a*, and *Usp18*, together with *Axl* and *Psmb9* (Fig 7C), but lacked broader induction of costimulatory or antigen-presenting machinery. These cells expressed high levels of phagocytic receptors characteristic of both M1 (*Fcgr3*, *Fcgr1*, and *Marco*) and M2 (*Mrc1* and *Cd163*) profiles without significant upregulation of downstream NF-κB- or STAT1-dependent inflammatory cascades, indicative of a basal or homing state. Notably, Cluster-3 was enriched in all three C1q subunits (*C1qa*, *C1qb*, *C1qc*), *C3*, *CtsK*, and *Nfatc1* (Fig 7C).

Cluster-4 macrophages were marked by high expression of *Stat6*, *Siglec1*, and *Cd40* (Fig 7C), in the absence of antigen-presentation machinery. This subset co-expressed high levels of *Msr1* and *Tnf*, alongside remarkably high levels of complement components *C1qa*, *C1qb*, and *C1qc* (Fig 7C), a signature aligned with the recently described M3-like complement-producing macrophage phenotype (57).

Finally, the identity of the osteoclast cluster was confirmed by the prominent expression of bone-remodeling machinery, including *Acp5*, *Ctsk*, *Nfatc1*, and *Mmp9* (Fig 7C). Furthermore, osteoclasts displayed robust expression of JAK-STAT components (*Stat3*, *Stat4*, *Stat5a*, *Stat5b*, and *Socs1*) and immune-modulatory markers (*Siglec1*, *Cd80*, *H2-K1*, and *Ciita*), pointing to a potential, albeit secondary, role in antigen presentation. Although phagocytic receptors were generally low (except *Cd209a*), osteoclasts co-expressed pro-inflammatory drivers (*Nfkb1*, *Il1b*, *Ccl2*, *Ccl5*, *Cxcl9*, *Cxcl10*, and *Nos2*) together with regulatory and tissue-repair factors (*Il10*, *Tgfb1*, and *Arg1*). These findings position joint-resident osteoclasts as key mediators bridging bone resorption, inflammation, and tissue homeostasis.

### 3.9. Persistent Viral RNA Sensing Drives IFN-π/IL-27–STAT1–Dependent Antiviral and Inflammatory Programs Across Joint-Associated Macrophage Subsets

In line with our hypothesis that MDM-CHIKV contribute to a chronic inflammatory state driven by persistent activation of viral RNA sensors and IFN-π/IL-27 pathway induction, Cluster-1 macrophages exhibited robust upregulation of key PRRs involved in viral sensing, including *Tlr2*, *Tlr3*, *Tlr7*, *Tlr8*, *Ifih1*/Mda5, *Dhx58*/Lgp2, and *Ifit1/2* (Fig 8A). In contrast, *Tlr4*, *Ddx58*/Rig-I, *Ddx60*, and *Eif2ak2*/Pkr were downregulated (Fig 8A). This selective pattern suggests sustained activation of specific viral RNA sensing in this subset, consistent with chronic viral RNA persistence (Fig 7B). Interestingly, Cluster-2 macrophages also upregulated *Tlr2*, *Tlr3*, *Ifih1*, *Ddx60*, and *Eif2ak2* (Fig 8A), further pointing to the selective activation of viral RNA sensors in alignment with persistent CHIKV RNA replication within this subset (Fig 7B). In contrast, Cluster-3 macrophages selectively upregulated *Tlr4*, *Ifih1*, and *Ifit2*, while downregulating *Tlr2* and *Tlr7* (Fig 8A). Cluster-4 macrophages displayed induction of *Tlr2* and *Ifih1*, accompanied by the downregulation of *Ddx60* (Fig 8A). Finally, osteoclasts showed no significant modulation of PRRs involved in viral recognition (Fig. 8A), suggesting that they do not actively participate in sensing CHIKV PAMPs in the joint microenvironment of chronically infected mice.

**Figure 8.**
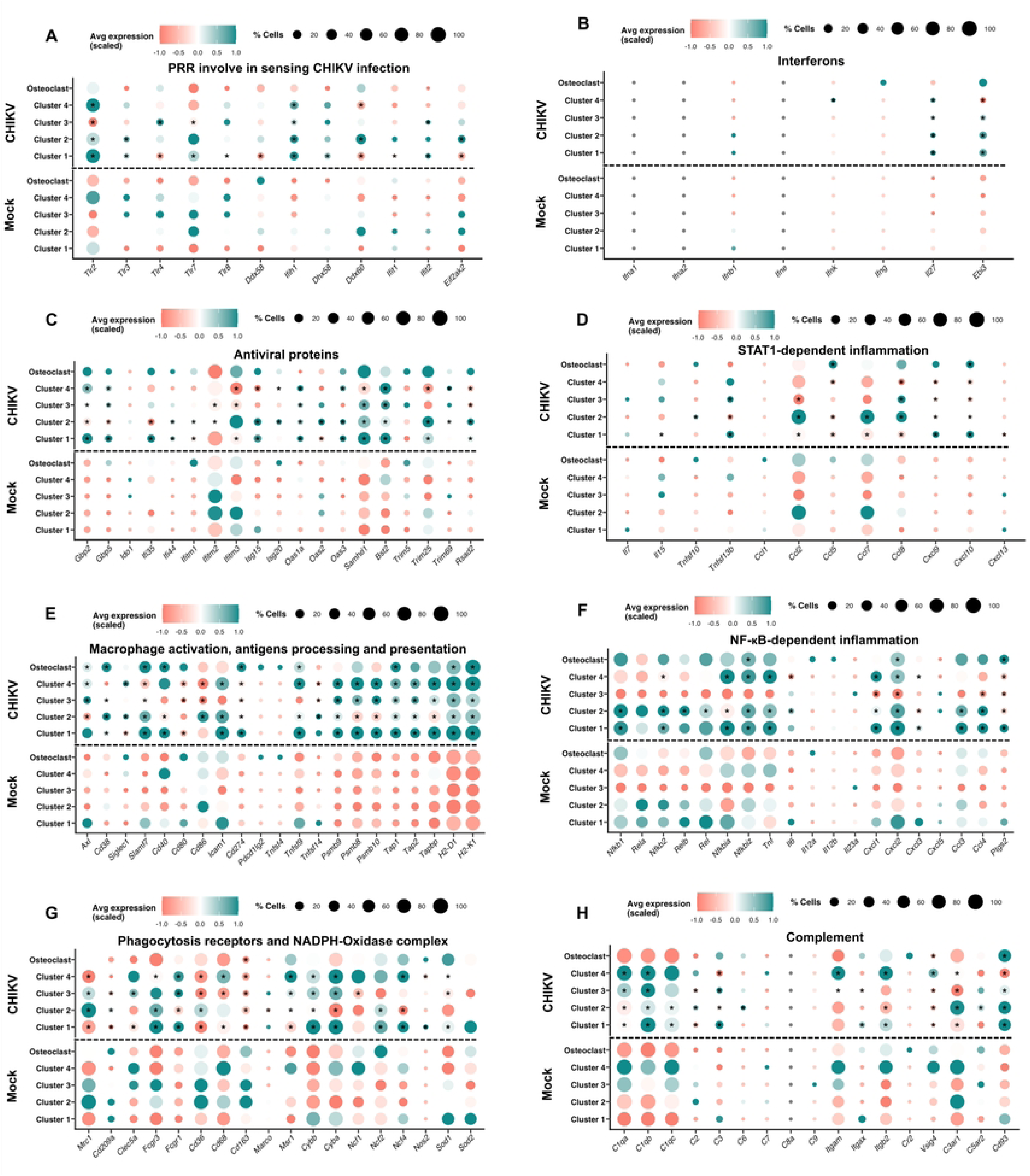
Persistent viral RNA sensing and multitiered NF-κB activation drive IFN-π/IL-27–STAT1–dependent inflammation, reprogrammed phagocytic machinery, and complement-mediated immunopathology in joint-associated macrophages during chronic CHIKV arthritis. (A-H) Dot plots illustrating expression profiles of key immune pathways across joint-associated macrophage subsets (Cluster 1-4 and Osteoclast) from mock and chronically infected joints (28 dpi): **A)** Pattern recognition receptors. **B)** Interferons and IL-27 subunits. **C)** Antiviral ISGs. **D)** STAT1-dependent inflammation genes. **E)** Antigen processing and presentation machinery. **F)** NF-κB-dependent inflammatory mediators. **G)** Phagocytic receptors and NADPH oxidase complex components. **H)** Classical, alternative, and terminal complement pathway components and receptors. Dot color reflects scaled average expression, and dot size indicates the percentage of expressing cells. Asterisks within dots denote statistically significant differential gene expression between CHIKV-infected and mock cells within each macrophage cluster, determined using a Wilcoxon rank-sum test via the FindMarkers function in Seurat. P-values were adjusted for multiple testing using the Benjamini-Hochberg method, with statistical significance defined at an adjusted p < 0.05.

Consistent with the chronic activation of PRRs involved in sensing CHIKV RNAs, we observed high induction of both IFN-π/IL-27 heterodimeric subunits (*Il27p28* and *Ebi3*) across joint-associated macrophage Clusters 1–3 in chronically infected mice, in the absence of canonical type I/II/III IFN induction (Fig 8B). This confirms the sustained *in vivo* expression of IFN-π/IL-27 during chronic CHIKV-induced arthritis. In contrast, cluster-4 macrophages uniquely expressed the *Il27p28* subunit while downregulating *Ebi3* (Fig 8B). Additionally, whereas *Ifnk* was specifically upregulated in cluster-4 macrophages, osteoclasts did not induce significant expression of any IFN gene (Fig 8B). Consistent with this cytokine profile, joint macrophages mounted a prominent antiviral response, wherein Clusters 1–4 expressed a broad spectrum of ISGs in a heterogeneous manner (Fig 8C). Notably, Cluster-1 and Cluster-2 macrophages from chronically infected mice displayed a significantly broader antiviral signature compared with other subsets, characterized by marked upregulation of *Ifi44, Isg15*, *Oas1a*, *Oas3*, *Samhd1*, *Bst2*/Tetherin, *Trim25*, and *Rsad2*/Viperin, aligning with their CHIKV-positive status and active engagement with viral RNA (Fig 7B). However, whereas Cluster-1 upregulated *Gbp2/5* and *Ifi35*, Cluster-2 downregulated these factors while specifically upregulating *Ifitm1*, *Isg20*, *Oas2*, and *Trim69* (Fig. 8C), revealing distinct antiviral transcriptional programs between two major infected macrophage subsets. Clusters 3 and 4 mounted a more restricted response, featuring significant induction of *Gbp2/5*, *Oas1a*, and *Bst2*, along with transcriptional repression of *Ifitm3* and *Rsad2* (Fig 8C). Finally, osteoclasts showed no significant changes in ISG expression during chronic CHIKV infection (Fig 8C).

In parallel with the induction of an antiviral state, we observed a marked upregulation of STAT1-dependent inflammatory mediators in joint-associated macrophages from chronically infected mice. Whereas Cluster 1 specifically upregulated *Il15*, *Tnfsf13b*/Baff, *Cxcl9*, and *Cxcl10*, alongside the downregulation of *Ccl2*, *Ccl5*, *Ccl7*, and *Ccl8* (Fig 8D). Cluster-2 macrophages mounted the strongest inflammatory response, significantly upregulating *Tnfsf10*/Trail, *Ccl2*, *Ccl7*, *Ccl8*, *Cxcl9*, and *Cxcl10*, alongside *Tnfsf13b* downregulation (Fig 8D). These findings demonstrate that these viral RNA-positive subsets orchestrate a heightened IFN-π/IL-27–STAT1-driven inflammatory program. In contrast, Cluster 3 upregulated *Tnfsf13b* and *Ccl8* while suppressing *Ccl2*, *Cxcl9*, and *Cxcl10* (Fig 8D), reflecting a more restricted inflammatory response. Cluster-4 macrophages only downregulated *Ccl5*, *Ccl8*, *Cxcl9*, and *Cxcl10*, whereas osteoclasts selectively induced *Ccl5* and *Cxcl10* (Fig 8D), highlighting the attenuated inflammatory engagement of these latter cell types during chronic murine CHIKV infection.

Consistent with the induction of robust IFN-π/IL-27-STAT1-dependent pro-inflammatory and antiviral responses, we found significant upregulation of macrophage activation markers and costimulatory molecules across all joint-associated macrophage subsets. Cluster-1 macrophages induced prominent upregulation of *Cd38*, *Slamf7*, *Cd40, Icam1*, *Cd274*/PD-l1, and *Tnfsf9*/4-1BB-L (Fig 8E). Similarly, Cluster-2 upregulated *Cd38*, *Siglec1*, *Slamf7*, *Cd86*, *Icam1*, *Cd274*, and *Tnfsf9* (Fig 8E), indicating strong inflammatory activation and stimulatory potential in both viral RNA-positive subsets during chronic CHIKV infection. Conversely, Cluster-3 and Cluster-4 macrophages upregulated *Axl* but downregulated *Slamf7, Cd80*, *Cd86*, and *Cd274* (Fig 8E), pointing to limited T-cell stimulatory capacity. Interestingly, osteoclasts upregulated *Axl*, *Cd38*, *Slamf7*, *Cd40*, *Cd274*, and *Tnfsf9* (Fig 8E), demonstrating inflammatory activation within this specialized cell lineage.

Additionally, most macrophage subsets exhibited marked upregulation of antigen processing and presentation machinery. Clusters 1, 3, and 4 displayed the highest upregulation of 20S-immunoproteasome catalytic subunits (*Psmb8*, *Psmb9*, and *Psmb10*), peptide transporter components (*Tap1*, *Tap2*, *Tapbp*), and MHC class I molecules, including *H2-D1* and *H2-K1* (Fig 8E), suggesting elevated capacity for antigen processing and presentation. Osteoclasts displayed a more restricted response, upregulating only *Tap1*, *H2-D1*, and *H2-K1* (Fig 8E). Crucially, Cluster 2 exhibited minimal or impaired expression of these antigen-presentation pathways, showing weak induction of *Psmb9*, *Tap1*, *Tap2*, *H2-D1*, and *H2-K1*, alongside significant downregulation of *Psmb8*, *Psmb10*, and *Tapbp* (Fig 8E). This suppression of the antigen-processing and presentation machinery in Cluster 2 may represent a viral immune evasion strategy, facilitating persistent CHIKV infection and replication within this specific macrophage reservoir.

### 3.10. Viral RNA-Positive Macrophages Sustain Multitiered NF-κB Activation, Reprogrammed Phagocytic Machinery, and Complement-Mediated Immunopathology in Chronic CHIKV Arthritis

Consistent with our observations in human MDM-CHIKV, joint-associated macrophages from chronically CHIKV-infected mice exhibited marked NF-κB pathway activation and induction of its canonical transcriptional targets. NF-κB-driven inflammatory signaling, a pan-myeloid transcriptional response, was observed across all macrophage clusters and osteoclasts, with robust upregulation of *Cxcl2* and *Ptgs2* and increased *Cxcl1* mRNA in macrophage clusters (Fig 8F). Clusters 1, 2, and 4 exhibited significant upregulation of *Nfkb2*, *NFkbia*, *Nfkbiz*, *Tnf*, and *Cxcl3*, whereas viral RNA-positive macrophages (Clusters 1 and 2) additionally exhibited significant upregulation of *Nfkb1* and *Ccl3* (Fig 8F). Furthermore, Clusters 1, 2, and 3 showed significant upregulation of *Ccl4*, whereas Cluster 2 was the only subset expressing *Relb* and *Rel*. Osteoclasts specifically displayed high mRNA levels of *Nfkbiz*, and *Il6* transcripts were strictly restricted to Cluster-2 and Cluster-4 macrophages (Fig 8F). Together, these findings demonstrate that viral RNA-positive macrophages sustain robust, multitiered NF-κB activation that promotes chronic joint inflammation, whereas osteoclasts exhibit a more restricted response.

Regarding phagocytic and ROS-producing capacities, joint-associated macrophage subsets exhibited distinct and highly variable receptor modulation profiles following chronic CHIKV infection. Cluster-1 macrophages upregulated activating phagocytic receptors, including *Fcgr3*/Cd16 and *Fcgr1*/Cd64, while downregulating M2-associated phagocytic receptors such as *Mrc1*/Cd206, C*d209a*/DC-SIGN, *Clec5a, Cd36*, Cd68, *Cd163,* and *Msr1* (Fig 8G), consistent with an M1-like immunophenotype. Additionally, Cluster-1 macrophages upregulated key NADPH oxidase components, including the catalytic subunits *Cybb*/Nox2 and *Cyba*/p22-phox, the regulatory factors *Ncf2*/p67-phox and *Ncf4*/p40-phox, and *Sod1*, as well as the inducible nitric oxide synthase *Nos2*/iNOS (Fig 8G). These expression profiles confirm that Cluster-1 macrophages possess a heightened capacity to promote phagocytosis-dependent oxidative burst within the joints of chronically infected mice.

In contrast, Cluster-2 macrophages uniquely upregulated *Mrc1*, *Fcgr3/*Cd16, *Cd36*, *Msr1*, and *Ncf2*, while downregulating *Cd209a*, *Fcgr1*, *Cd163*, *Marco*, *Cyba*, and *Ncf4* (Fig. 8G). This profile is indicative of a predominant M2-like phenotype with enhanced C-type lectin/scavenger receptor-driven phagocytic programs, accompanied by a limited capacity for ROS production compared to Cluster-1 macrophages. Cluster-3 macrophages exhibited upregulation of *Mrc1*, *Clec5a*, *Fcgr1*, and *Msr1*, but downregulated a broad array of phagocytic receptors, including *Cd209a*, *Cd36*, *Cd68*, and *Cd163* (Fig 8G), suggesting a marked reduction in overall phagocytic capacity. However, Cluster 3 upregulated *Cybb* and *Cyba*, while downregulating *Ncf1* and *Sod1* (Fig 8G), pointing to a partial capacity for ROS generation during chronic CHIKV infection. Cluster-4 macrophages displayed increased expression of *Fcgr3*, *Fcgr1*, *Cd68*, and *Msr1*, alongside concurrent downregulating of *Mrc1*, *Cd36*, and *Cd163* (Fig 8G). Additionally, Cluster 4 upregulated *Cybb*, *Cyba*, and *Ncf4*, but downregulated *Nos2* and *Sod1*, likewise reflecting a partial ROS-producing potential. In turn, osteoclasts exhibited the most restricted transcriptomic alterations, showing a specific reduction only in *Cd163* expression (Fig 8G).

Finally, we evaluated complement system activation across joint-associated macrophage subsets in chronically infected mice. Consistent with our observations in human MDM-CHIKV, *C1qa* and *C1qb* were significantly upregulated across Clusters 1–4 (Fig 8H). However, Cluster-1 and Cluster-2 macrophages also upregulated C1qc, C3, and Cd93 (Fig. 8H), indicating a heightened capacity to initiate the classical complement pathway within these viral RNA-positive subsets. Regarding terminal complement components and receptors, Cluster-1 macrophages upregulated *Itgax*/Cd11c and *Itgb2*/Cd18 while downregulating *C2, Vsig4*, and *C3ar1* (Fig. 8H), suggesting an enhanced capacity to promote phagocytosis of complement-opsonized antigens. Cluster-2 macrophages displayed concurrent upregulation of *C2*, *C6*, *C3ar1*, and *C5ar2*, alongside downregulation of *Itgb2* and *Vsig4* (Fig 8H).

Cluster-4 macrophages showed a particularly striking phenotype, exhibiting high basal expression of C1q subunits even under mock conditions (Fig 8H), a hallmark of the recently defined complement-producing M3 macrophage subset (57), which was further elevated upon chronic CHIKV infection. Cluster-4 macrophages further upregulated *Itgam*/Cd11b, *Itgb2*, and *Vsig4*, but repressed *C3* and *Cd93* (Fig 8H), suggesting a specialized role in local complement secretion and complement-mediated phagocytosis. Finally, osteoclasts showed the fewest transcriptomic alterations in response to CHIKV infection, characterized solely by upregulation of *Cd9*3 and downregulation of *Vsig4* (Fig 8H).

Collectively, these findings reinforce our hypothesis that CHIKV actively redirects monocyte-to-macrophage differentiation toward a specialized, mixed M1-like activation state capable of sustaining viral replication over extended periods. Within the joint microenvironment during *in vivo* infection, these macrophages serve as the primary viral reservoir while establishing a chronic pro-inflammatory niche driven by synergistic IFN-π/IL-27–STAT1 and NF-κB signaling. Furthermore, their reprogrammed phagocytic machinery and heightened complement activation perpetuate local immunopathology and tissue destruction underlying chronic arthritis. Together, these mechanisms highlight joint-associated macrophages as a central pathogenic axis and a promising therapeutic target for mitigating persistent joint inflammation in chronic CHIKV disease.

## 4. Discussion

Our study provides comprehensive mechanistic insights into the role of monocytes and macrophages in CHIKV immunopathogenesis, with direct implications for chronic arthralgia. By integrating *ex vivo* transcriptomic signatures from acute and convalescent CHIKD patients, *in vitro* myeloid differentiation models, and *in vivo* single-cell RNA-seq profiling of joint-associated macrophages from chronically infected WT C57BL/6J mice, we uncover a complex interplay among viral sensing, macrophage reprogramming, and divergent inflammatory pathways that shape the transition from acute infection to persistent joint disease.

Our initial observation that pediatric CHIKD patients exhibit robust upregulation of monocytic lineage markers and phagocytic receptors in peripheral blood provides compelling evidence that monocyte-to-macrophage differentiation is actively engaged during natural infection. This expands upon previous reports describing shifts in monocyte subsets during CHIKV infection (23)(34,35)(37), establishing that viral exposure directly drives monocyte commitment and functional reprogramming rather than merely infecting pre-existing myeloid pools. The simultaneous induction of multiple PRR families, including TLRs, RLRs, and inflammasome components, confirms that CHIKV sensing engages broad innate surveillance machinery (37)(58,59)(60). Notably, the most striking *in vivo* observation was sustained, multiple interferon responses persisting from acute infection into convalescence, coupled with continuous JAK-STAT activation and an expansive ISG signature encompassing myeloid-restricted effectors such as APOBEC3A and IDO1 (11). This underscores that monocytes rely on IFN-triggered cell-intrinsic mechanisms to achieve antiviral resistance (Fig. 9).

**Figure 9.**
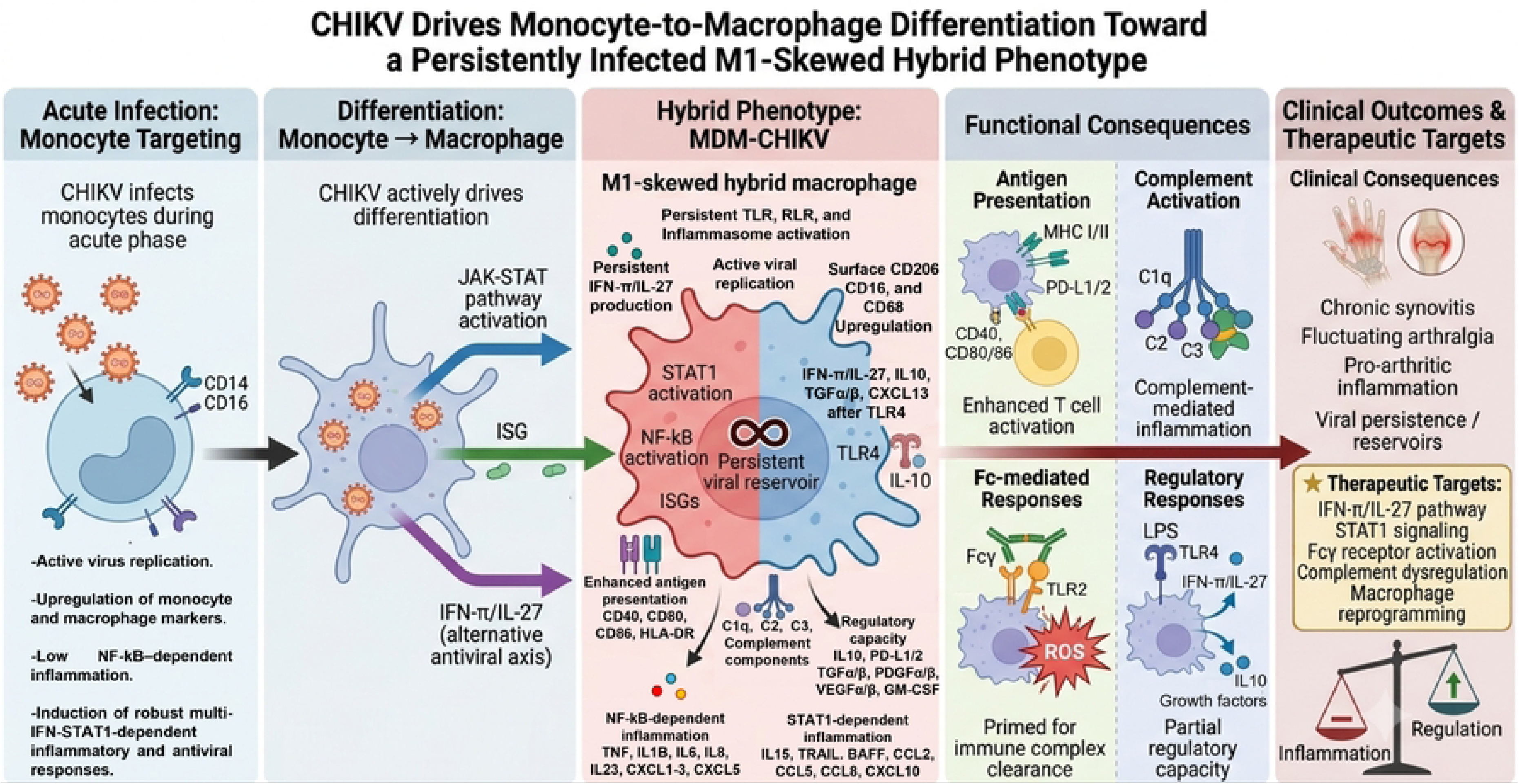
Proposed mechanism of CHIKV-driven monocyte-to-macrophage differentiation and chronic arthritogenic inflammation. Schematic representation summarizing how CHIKV infection during monocyte-to-macrophage differentiation established a persistently infected, M1-skewed hybrid effector phenotype. This differentiation program fuels viral persistence, drives IFN-π/IL-27-STAT1 and NF-κB signaling, and reprograms phagocytic and complement machinery, thereby sustaining immunopathology and tissue remodeling in chronic CHIKV-induced arthritis.

The *in vitro* differentiation model recapitulated and extended these findings, revealing that CHIKV infection during monocyte-to-macrophage differentiation drives persistent viral replication accompanied by profound morphological and transcriptomic remodeling. MDM-CHIKV cells maintained active viral RNA replication through day 6 post-infection despite an initial robust ISG response (23)(37), indicating that macrophages may serve as cellular viral reservoirs, continuously fueling antigen presentation and secretome activation, thereby driving persistent tissue inflammation (32). The observed morphological shifts, an elongated, ameboid phenotype with reduced cell size and complexity, likely reflect virus-induced cytoskeletal reorganization that may alter macrophage migration and intercellular communication within the synovial microenvironment (61).

Our scRNA-seq analysis of joint-associated macrophages from chronically CHIKV-infected WT C57BL/6J mice provided critical *in vivo* validation of these findings. We identified five transcriptionally distinct macrophage clusters, with cluster 1 and cluster 2 representing the predominant subsets (together ∼62% of the population). Crucially, while both CHIKV nsP- and sP-ORFs were detected in cluster-1 and cluster-2 macrophages, the viral RNA (−) replicative intermediate was exclusively found in cluster-2 macrophages, confirming that this subset constitutes the bona fide replicative viral reservoir within the arthritic joint. This observation directly supports our hypothesis that persistently infected macrophages drive chronic inflammation, and it establishes cluster-2 macrophages as the primary cellular sanctuary for CHIKV *in vivo*.

Comparative transcriptomic analysis positioned MDM-CHIKV cells closest to classically activated M1 macrophages, sharing hallmark features such as STAT1- and NF-κB-dependent inflammatory programs, robust ISG induction, and upregulation of antigen presentation machinery (Fig. 9). However, MDM-CHIKV cells also displayed unique transcriptomic features that distinguished them from canonical M1 macrophages. The selective induction of IFN-π/IL-27, rather than canonical type I/III IFNs during late differentiation, represents a key divergence with major implications for chronic pathology. IFN-π/IL-27 is a pleiotropic cytokine with both pro-inflammatory and immunoregulatory properties; its sustained production may contribute to chronic arthralgia by driving STAT1-dependent chemokine cascades while failing to achieve complete viral clearance (6)(13)(11,14). Crucially, in joint-associated macrophages from chronically infected mice, we observed significant induction of both *Il27p28* and *Ebi3* across clusters 1–3, confirming chronic IFN-π/IL-27 production *in vivo*. Moreover, cluster-1 and cluster-2 macrophages exhibited the most robust ISG response and highest expression of STAT1-dependent chemokines (*Ccl8*, *Cxcl9*, *Cxcl10*), alongside NF-κB activation and upregulation of pro-inflammatory targets (*Tnf*, *Cxcl1–3*, *Ccl3*, *Ccl4*, *Ptgs2*/Cox2). These findings establish that the virus-permissive macrophage subsets are also the principal drivers of IFN-π/IL-27–STAT1 and NF-κB-mediated inflammation within the joint (Fig 9).

The divergent activation dynamics between NF-κB and STAT1 pathways indicate fundamental reprogramming of inflammatory signaling in CHIKV-exposed myeloid cells. While NF-κB-dependent cytokines such as TNF-α and IL-6 were modestly induced, STAT1-dependent chemokines including CCL-2, CCL-5, and CXCL-10 were markedly upregulated. This chemokine-biased output is directly relevant to CHIKV-associated arthropathy, as these chemo-attractants recruit monocytes, T cells, B cells, and NK cells into synovial tissues, perpetuating localized joint inflammation (6,7)(30)(62)(63). Notably, persistent CXCL-10 is a hallmark of rheumatoid arthritis and other chronic autoimmune arthropathies, underscoring shared pathogenic mechanisms (63)(64).

Functional characterization of MDM-CHIKV cells revealed additional regulatory complexity. Despite displaying an M1-like global signature, these cells exhibited surface upregulation of M2-associated markers including CD206 and demonstrated a distinct, context-dependent phagocytic capacity. Their attenuated ROS production in response to unopsonized zymosan versus amplified respiratory burst upon serum-opsonized challenge indicates that CHIKV-exposed macrophages are primed for opsono-phagocytic clearance. This heightened oxidative capacity, while advantageous for clearing opsonized virions, may also drive FcγR-mediated inflammatory activation in the presence of antiviral or cross-reactive autoantibodies, promoting antibody-dependent enhancement of pathology and persistent synovial inflammation (Fig 9) (65)(66)(67). Consistent with this, our scRNA-seq data revealed variable phagocytic receptor modulation across clusters, suggesting a complex, subset-specific functional polarization.

MDM-CHIKV cells also exhibited a pronounced complement activation gene signature, including upregulation of *C1q*, *C2*, and *C3* components, indicating capacity for local synthesis and activation of both classical and alternative pathways (Fig 8) (68)(69). Our scRNA-seq analysis confirmed significant upregulation of *C1qa* and *C1qb* across clusters 1– 4, with cluster 1 and cluster 2 additionally upregulating *C1qc*, *C1r*, and *C1s*, suggesting stronger complement activation in the virus-permissive subsets. Notably, cluster-4 macrophages displayed high basal *C1q* expression even under mock conditions, a phenotype associated with the recently described M3 profile of macrophages with high complement-producing capacity (57). While complement-mediated opsonization facilitates viral clearance, unconstrained complement activation triggers collateral tissue injury through membrane attack complex assembly and localized anaphylatoxin release (Fig 9) (53)(70). The downregulation of complement regulatory receptors in certain clusters may further disrupt homeostasis, shifting the microenvironment toward pro-inflammatory outcomes.

Perhaps most critically for chronic arthralgia pathogenesis, MDM-CHIKV cells displayed markedly enhanced antigen-presentation capacity, characterized by upregulation of immune-proteasome subunits, TAP transporters, MHC class I/II molecules, and costimulatory ligands (Fig 9). This machinery endows CHIKV-exposed macrophages with the capacity to vigorously stimulate both CD8+ and CD4+ T-cell subsets. While a robust T-cell response is essential for acute viral clearance, persistent antigen presentation during ongoing replication may induce T-cell exhaustion or promote autoreactive responses through bystander activation or molecular mimicry (71)(72). The upregulation of PD-L1 and PD-L2 alongside CD80/CD86 suggests complex regulation of T-cell responses, potentially contributing to the balance between protective immunity and immunopathology (71)(72).

Upon secondary challenge with LPS, MDM-CHIKV cells upregulated M2-associated regulatory factors including *IL10*, *TGFA*, and *TGFB1*, demonstrating retained capacity for anti-inflammatory responses (25)(18,73). However, concurrent IFN-π/IL-27 production, sustained STAT1-dependent chemokine expression, and maintained antigen presentation indicate that this regulatory response is insufficient to fully resolve inflammation. This hybrid immunophenotype, simultaneously expressing pro-inflammatory, antiviral, regulatory, and tissue-remodeling programs, provides a compelling mechanistic framework for the relapsing-remitting course of post-CHIKV arthralgia, in which joint symptoms wax and wane in response to intercurrent infections or environmental triggers (6)(4,8).

Collectively, our findings support an integrative model in which CHIKV infection of monocytes drives their differentiation into persistently infected macrophages with an M1-skewed, hybrid phenotype. These cells establish long-term viral reservoirs, particularly cluster-2 macrophages within the joint, produce IFN-π/IL-27 alongside STAT1-dependent inflammatory effectors, activate NF-kB, complement and opsono-phagocytic machinery, and expand antigen presentation capacity while retaining partial immunoregulatory features. The resulting microenvironment, sustained by chemokine-driven leukocyte recruitment, complement activation, and persistent T-cell stimulation, creates a permissive niche for chronic synovitis and arthralgia (Fig 9). These insights position maladaptive macrophage reprogramming as a primary determinant of chronic sequelae and support therapeutic strategies targeting macrophages, IFN-π/IL-27–STAT1 signaling, complement dysregulation, or FcγR-driven macrophage activation to prevent irreversible joint injury.

## 5. Limitations and perspectives

While this study provides high-resolution insights into the immunopathogenesis of chronic CHIKV-induced arthritis, several limitations warrant consideration. First, our study did not include the quantification of MDM-CHIKV expressing the CHIKV-E antigen; therefore, it was not determined what proportion of MDM were infected and capable of active replication. Second, although the C57BL/6J mouse model accurately recapitulates key features of alphaviral synovitis, human synovial architecture and immune cell kinetics may exhibit distinct heterogeneity; thus, translational validation in clinical synovial biopsies is required. Additionally, our analysis focused on a single chronic time point (28 dpi); future longitudinal fate-mapping studies will be essential to establish the exact longevity and turnover of viral RNA-positive macrophage reservoirs. Finally, while scRNA-seq revealed key signaling networks, including the IL-27–STAT1 and multitiered NF-κB axes, further functional studies utilizing lineage-specific knockout models will be required to definitively prove the causative roles of individual macrophage subsets in driving long-term joint pathology.

### Use of Artificial Intelligence Tools statement

Artificial intelligence (AI)-assisted tools were used in a limited capacity during the preparation of this manuscript. Specifically, Claude and DeepSeek were used to assist in improving the clarity, grammar, and overall readability of the text, while Grammarly was used for language editing. These tools were employed solely to refine the expression of the authors’ original writing and did not contribute to the generation or interpretation of scientific content. In addition, DeepSeek was used to assist in generating an initial prompt for the conceptual design of the graphical model presented in Fig 2A and Fig 9. This prompt was subsequently used in the generative AI system Gemini to produce a preliminary visual backbone of the graphical summary. The resulting output served only as an initial conceptual template and was extensively modified, curated, and finalized manually by the authors using PowerPoint to ensure scientific accuracy, appropriate representation of the underlying biological concepts, and consistency with the data presented in the manuscript. All scientific ideas, experimental design, data analysis, interpretations, and conclusions are the sole responsibility of the authors.

